# Understanding the complex interplay between tau, amyloid and the network in the spatiotemporal progression of Alzheimer’s Disease

**DOI:** 10.1101/2024.03.05.583407

**Authors:** Ashish Raj, Justin Torok, Kamalini Ranasinghe

## Abstract

**INTRODUCTION:** The interaction of amyloid and tau in neurodegenerative diseases is a central feature of AD pathophysiology. While experimental studies point to various interaction mechanisms, their causal direction and mode (local, remote or network-mediated) remain unknown in human subjects. The aim of this study was to compare mathematical reaction-diffusion models encoding distinct cross-species couplings to identify which interactions were key to model success.

**METHODS:** We tested competing mathematical models of network spread, aggregation, and amyloid-tau interactions on publicly available data from ADNI.

**RESULTS:** Although network spread models captured the spatiotemporal evolution of tau and amyloid in human subjects, the model including a one-way amyloid-to-tau aggregation interaction performed best.

**DISCUSSION:** This mathematical exposition of the “pas de deux” of co-evolving proteins provides quantitative, whole-brain support to the concept of amyloid-facilitated-tauopathy rather than the classic amyloid-cascade or pure-tau hypotheses, and helps explain certain known but poorly understood aspects of AD.

## BACKGROUND

Alzheimer’s disease (AD) involves widespread and progressive deposition of amyloid beta (Aβ) protein in cortical plaques and tau tangles^1,2^. Aβ usually first appears in frontal regions and subsequently spreads to allocortical, diencephalic, brainstem, striatal and basal forebrain regions ^2,3^. In contrast to Aβ, tau tangles appear first in the locus coeruleus, then entorhinal cortex, followed by an orderly spread into hippocampus, amygdala, temporal lobe, basal forebrain, and isocortical association areas^1^. The dominant “amyloid cascade hypothesis” ^4^, posits that amyloid is the upstream factor, whose early abnormal accumulation in the brain causes a cascade of downstream events that recruit misfolded tau. However, this hypothesis has encountered several difficulties, including the spectacular failure of many large clinical trials of amyloid-targeting therapies, and the fact that the temporal and regional distribution of Aβ is quite dissociated from that of tau as well as of downstream atrophy and cognitive deficits^5–7^. This has led to the search for other mechanisms, especially the role of tau, which is increasingly considered more central to AD pathophysiology. However, it is unlikely that tau alone can explain AD pathogenesis, due to overwhelming mechanistic, genetic and demographic evidence of amyloid involvement.

Among potential alternative concepts put forward to reconcile these difficulties, a prominent one is the emerging concept of network-based transmission of both amyloid and tau. Evidence is accumulating in favor of self-assembly and trans-neuronal propagation of amyloid and tau, indeed, of all neurodegenerative pathologies ^8–11^. Unlike conventional assumptions about spatial spread, this emerging concept implies spread along axonal projections, and the resulting networked spread has repeatedly been substantiated by neuroimaging ^12–15^. Network-mediated spread of amyloid and tau offers an attractive means of reconciling above difficulties: the two species may interact locally, but the effects of these interactions do not remain local, and may propagate across neural circuits to distant regions. Given that the two species have unique and different “epicenters” or seeding loci, this would lead to the apparent observation of their deposition in separate non-overlapping regions. This broad hypothesis as schematized in **Figure 1A**, posits that the spatiotemporal progression of AD-related amyloid and tau proceeds on the connectivity network after being seeded at different sites. As the progression proceeds, we expect that the two entities will come in contact with each other, interact kinetically by affecting the other species aggregation and spread. Thus, the cross-species interaction, in the background of network-mediated propagation, may present a plausible framework for understanding AD progression.

**Figure 1.**
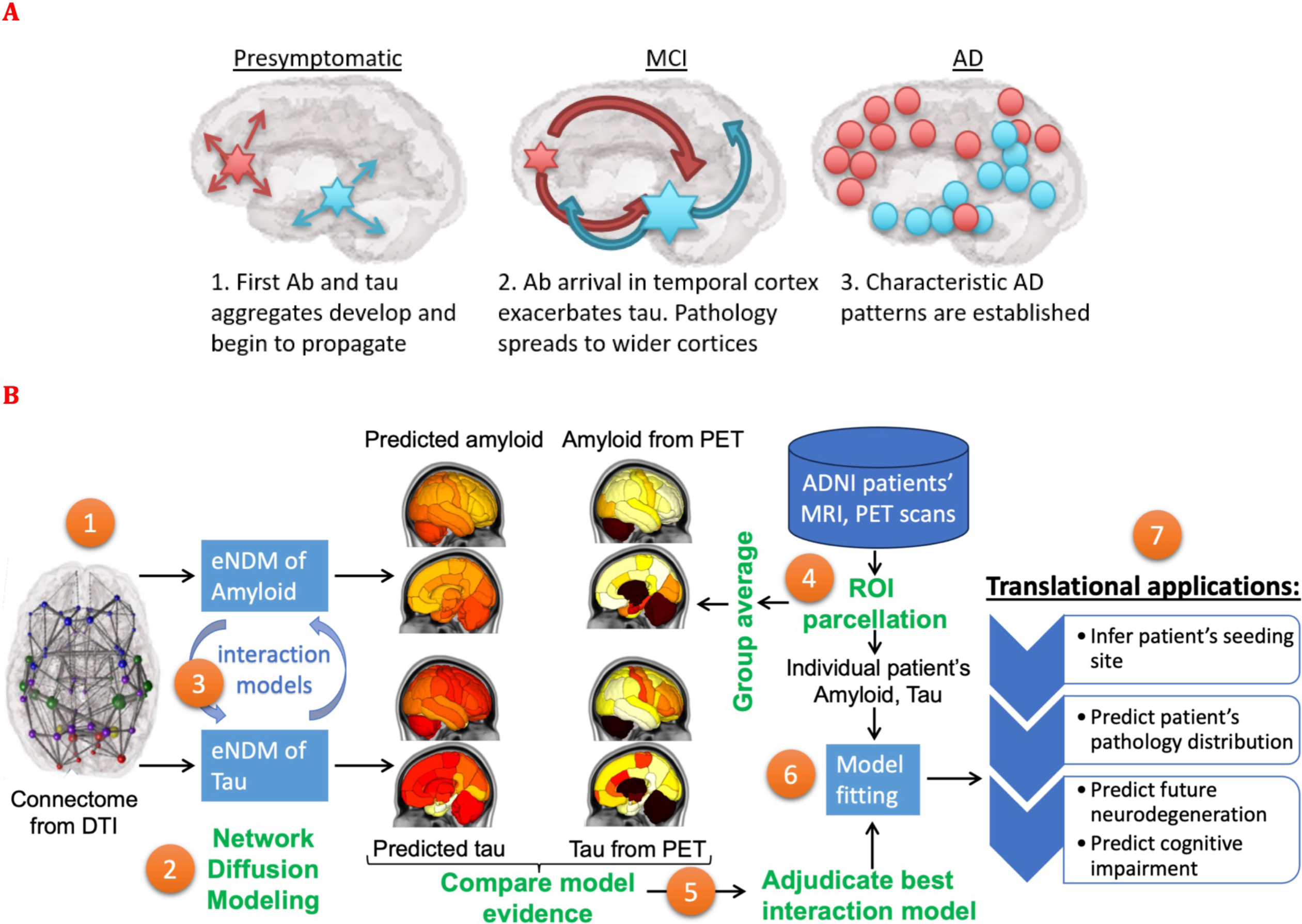
Study goals, design and work flow. **A**: This study aims to test the broad hypothesis schematized here, such that the spatiotemporal progression of AD-related amyloid and tau proceeds on the connectivity network after being seeded at different sites. As the progression proceeds, we expect that the two entities will come in contact with each other, interact kinetically by affecting the other species aggregation and spread. **B**: The overall study design to test these processes is illustrated here. Using the structural network connectome (**1**), we deployed the extended Network Diffusion Model (eNDM) that seeks to recapitulate trans-neuronal spread of amyloid and tau pathology (**2**). Starting with the base model of independent eNDMs for both amyloid and tau, we successively introduced various models of interaction between the two (**3**) – these models are listed in **Table 1**. From the ADNI database we extracted regional distributions of tau and amyloid SUVr from raw PET images (**4**), whose group average patterns were used to statistically evaluate against each computational model (**5**). Using model evidence criteria we adjudicated between all possible interaction models and selected the best performing one, yielding a numerical assessment of the most likely mode of interaction between amyloid and tau. This adjudicated model was then fit to individual patients’ regional pathology data using Bayesian inference of model parameters (**6**). The fitted model was capable of not only predicting the patient’s pathology pattern but also their specific seeding sites – a potential marker of translational interest. The model outcomes may in future translational applications be used to predict future pathology, neurodegeneration and cognitive state of the patient (**7**).

**Table 1:**
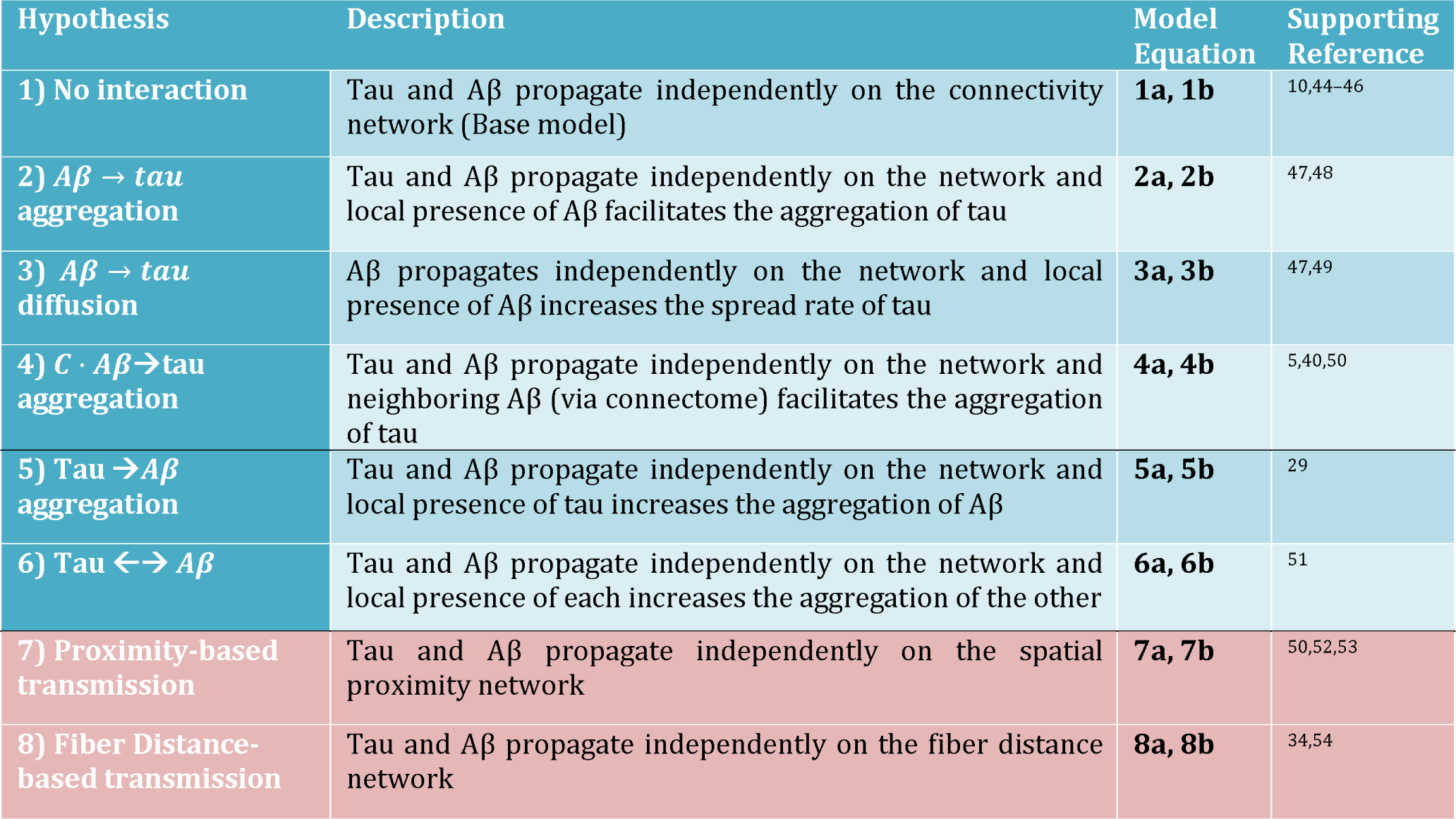
Alternative/competing hypotheses of tau-amyloid interaction to be tested in the background of transmission mediated by network connectivity. Brief description of each hypothesis is given, along with the mathematical differential equation corresponding to it, contained in **SI:Note 1**. References to supporting literature for each hypothesis is provided. Two additional models of spread are given, pertaining to spread via proximity and spread via fiber distance – for these models we use the base case of no interaction between tau and amyloid.

Many experimental and mechanistic studies are available that point to a complex interaction between amyloid and tau. It is well known that Aβ facilitates the aggregation of tau and influences the course and severity of downstream atrophy^16–18^. Mechanistic studies point to various modes of interaction, both direct and indirect^17,19–21^. Hippocampal injection of AD-derived tau into the brains of transgenic mice expression mutant APP was found to promote tau aggregation in “dystrophic neurites”, which have an Aβ core, as well as neurofibrillary tangles and neuropil threads, suggesting a direct, local interaction. ^22^ These different species were found to accumulate and spread at different rates as function of Aβ plaque load^23^. Alternatively, others have proposed that Aβ mediates the accumulation and spread of tau by recruiting microglia and inducing a pro-inflammatory immune response^24–27^, or by causing hyperexcitability-mediated release of tau^28^. Tau may also play a role in Aβ formation, which may occur in the absence of Aβ-mediated tau accumulation^29^ or with the two species interacting synergistically^30^.

While these studies provide important mechanistic evidence in model organisms, they may be sensitive to idiosyncratic methodological choices, leading to a diversity of mechanistic possibilities that may not generalize and may not be germane to human disease. Key aspects of brain-wide propagation of tau and amyloid, and the exact mechanistic form of their mutual interactions, therefore remain unsupported empirically in human AD. Since experimental testing of mechanistic hypotheses is difficult in humans, here we took a somewhat unorthodox approach, by relying on a *computational* rather than *experimental* interrogation of the mechanistic interaction between tau and amyloid. Our overall study design to test the above processes is illustrated in **Figure 1B**.We first formulated a mathematical model that recapitulates the spatiotemporal evolution of human AD, based on recent advances in the mathematical modeling of reaction-diffusion or network processes that have emerged as a powerful means of evaluating the brain-wide consequences of biophysical mechanisms underlying self-assembly and propagation of neurodegenerative pathologies ^14,15,31–36^. On top of this base network model, we then built various *interaction models* that allow Aβ and tau species to interact in local neural populations. We compared the performance of each interaction model via thorough statistical adjudication, by accumulating model evidence on large neuroimaging (tau and amyloid PET, MRI) datasets of AD spectrum subjects. We devised a fitting procedure to obtain model parameters that best match individual subjects’ regional disease patterns. In this manner we were not only able to determine the most well-supported modes of cross-species interactions, but also their applicable kinetic rates. Although spatial and network spread models of single pathological species have been previously reported^14,15,31–36^, and data-driven models of multiple biomarkers are also available^36–38^, this study is unique in modeling and evaluating directly on empirical data the network-mediated transmission and interaction of tau and amyloid *jointly*.

We show that both network propagation and amyloid-tau interaction are necessary to recapitulate human AD data. Our data conclusively support that network-mediated spread with a 1-way interaction, whereby amyloid facilitates local tau aggregation, is the most parsimonious and accurate model, yielding correlations above 0.7 against empirical tau topography. We also tested other interactions, including bidirectional ones and those involving enhanced pathology spread instead of aggregation, but these were not well supported on statistical tests. In totality, this “toxic *pas de deux*”^17^ of co-evolving tau and amyloid pathologies provides critical numerical support to mechanistic hypotheses not possible to be tested directly in humans. We anticipate that our computational testbed will become an important future tool for the generation and testing of mechanistic hypotheses with the potential to complement studies in model organisms.

## MATERIALS AND METHODS

### Experimental Design

#### Subjects and data

Data used in this study were obtained from the ADNI ^41^ database (http://adni.loni.usc.edu); consisting of 531 ADNI-3 subjects who had at least one exam of all three: MRI, AV1451-PET and AV45-PET, available by 1/1/2021. Demographic information is in **Table S1.** These data were processed to obtain regional of pathology and atrophy, the latter used in this analysis as a measure of tau-induced neurodegeneration. Anatomic connectomes were computed from healthy diffusion MRI and tractography algorithms. The primary dataset was evaluated on the 86-region Desikan atlas and the Supplementary dataset on 90-region AAL atlas, using similar processing pipelines; see **SI: Note 2.** To remove AV1451-PET scans’ non-specific binding and the effect of iron in thalamus and striatum, their values were removed from subsequent analysis, leaving 76 regions. Proposed network models were applied to canonical healthy connectomes from human connectome project (HCP)^42^ and model patterns compared against the ADNI regional data.

### Network spread model with cross species interactions

Refer to **SI: Note 1** for detailed model description. Here we summarize the overall amyloid-tau coupled system:

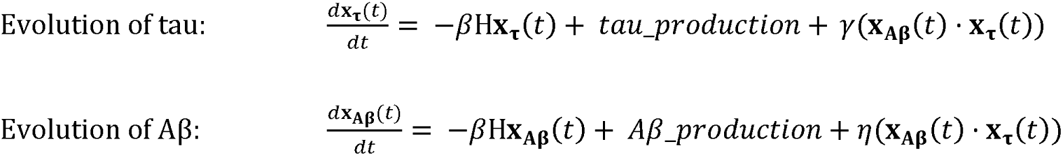

The first term on the right represents network diffusion, following our prior work ^12^, whereby pathology spread follows regional concentration gradients restricted along network connections. This involves the connectome’s Laplacian matrix H and the diffusivity rate constant β. This model captures trans-neuronal propagation as a connectivity-based process. The second term specifies how pathologic tau and amyloid respectively are initially seeded and produced during the course of progression (**Figures S1** and **S2**). The last term encodes the interaction between the two species, and is the object of specific interest in this study. Accordingly, from the broad system above we have derived several network-interaction models of increasing complexity (Eqns 1-8 in **SI)** and carefully tested them against each other.

### Statistical Analysis and model testing

Both group statistics (t-statistics for each of EMCI, LMCI and AD groups) and individual fitting was performed. Empirical regional AV1451-PET tau was supplemented with MRI-derived regional atrophy to leverage larger sample size; since atrophy is a useful surrogate for tau, with strong regional association ^43^. The statistical test of choice is Pearson correlation strength, R, and its two-tailed p. Detailed model fitting to individual subjects is described in **SI: Note 6.** We used a Bonferroni correction to account for potential false positives. We also performed extensive permutation tests with 500 random permutations and compiled additional measures of significance. We used Fisher’s R-to-z transformation to assess significance between models of comparable complexity, and AIC for models with varying complexity. **Data and code Availability.** Patient data can be directly obtained from the ADNI study (http://adni.loni.usc.edu). To facilitate review, group data herein will be made available publicly and without limitations, along with the entire code repository, at our laboratory’s GitHub site: https://github.com/Raj-Lab-UCSF/Aggregation-Network-Diffusion. There are no restrictions or embargoes, subject to standard BSD3 license.

## RESULTS

### I. Aggregate relationships between tau, amyloid, atrophy and the network

The public ADNI3 data^41^ was processed with established software pipelines to obtain regional imaging biomarkers of atrophy, tau and amyloid (demographics in **Table S1)**. First, we ascertained how individual subjects’ biomarker triplets are related to each other at the aggregate level. We find a moderate yet significant (p < 10^-3^, post-Bonferroni correction) relationship between atrophy and tau, but not between atrophy and amyloid (**Figure 2A**). There is a strong relationship between tau and amyloid, providing an empirical justification for modeling local amyloid-tau interactions^16–18^. Accompanying histograms of correlation strengths of disease groups (EMCI, LMCI, AD) demonstrate a prominent stage dependence, whereby atrophy is more tightly related to tau in later rather than earlier stages. The situation is reversed for tau-amyloid associations: earlier stages have a stronger association than later stages. These data point to the well-known finding that amyloid plays a role early in AD pathophysiology, and at later stages it has a plateauing behavior and is no longer predictive. **Stage-dependent relationship to network connectivity.** To assess the hypothesis that biomarkers are predicted by connectivity to pathology origination site at the aggregate level, we plotted a region’s biomarkers against its network connectivity to bilateral EC (**Figure 2B)**. We find moderate but significant (p < 10^-3^, post-Bonferroni) association between EC-connectivity and all three biomarkers; however, the association with amyloid is in fact negative. These results are also stage-dependent; with earlier stages giving a stronger association with connectivity for atrophy and tau, and the reverse for amyloid. **Longitudinal relationships.** The longitudinal change (difference between baseline and year-1 visit) of biomarkers was plotted against baseline in **Figure 2C**. A moderate but significant association (p < 10^-3^, post-Bonferroni) with change of tau was found for baseline tau, but not baseline amyloid. Next we broadly assess network involvement and potential remote effect, i.e. whether baseline pattern of tau weighted by network connectivity would predict the change of tau. We used a simple model of spread of tau from baseline pattern along the network, given by the well-established network diffusion model (Raj 2012). The hypothesis appears to have significant support for baseline tau, but not for amyloid, and incurs a stage dependency as with earlier results. *Taken together, these results demonstrate significant and stage-dependent cross-sectional and longitudinal relationships between tau and amyloid distributions in the Alzheimer brain*.

**Figure 2.**
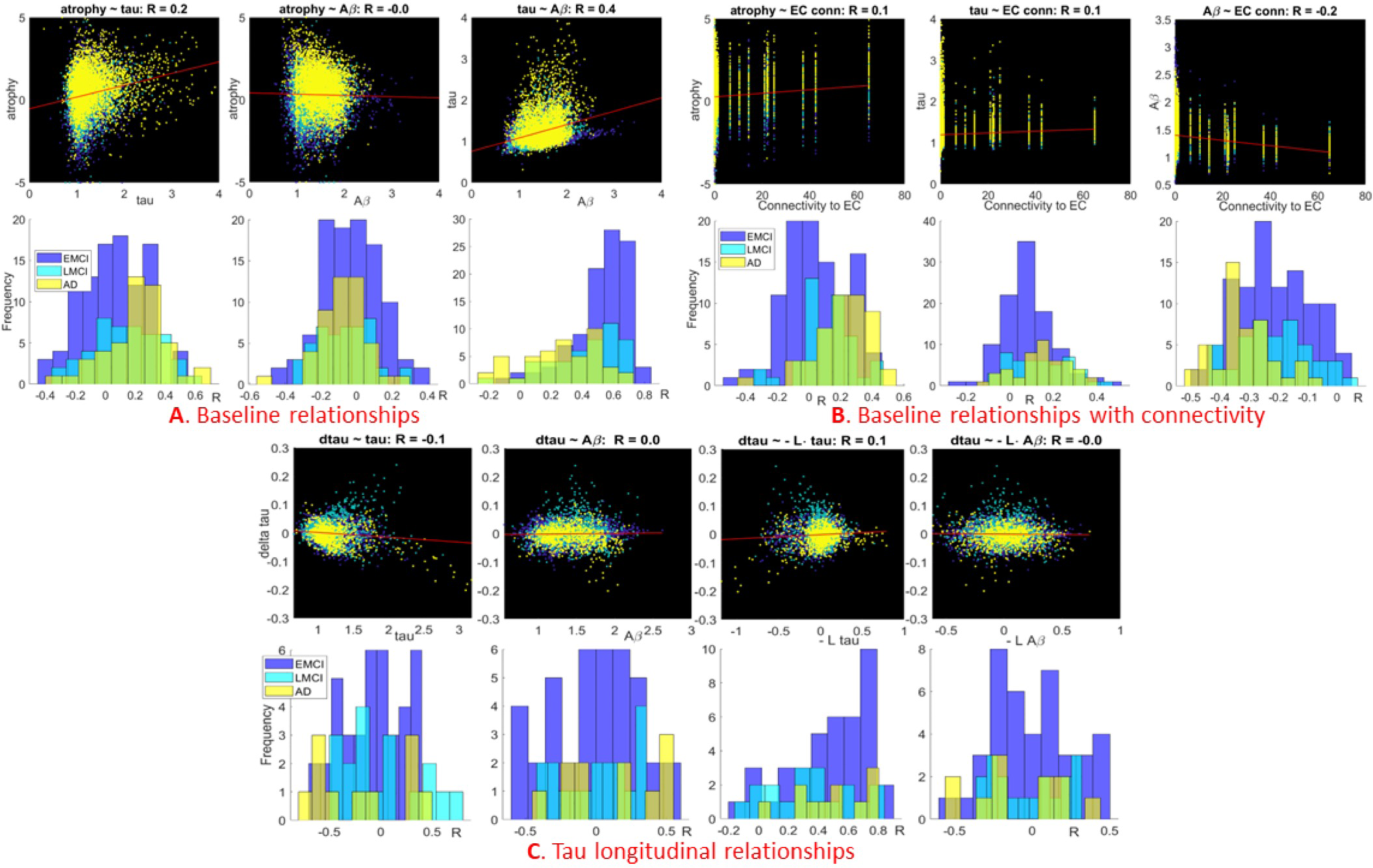
Cross-sectional and longitudinal relationships between tau, amyloid, atrophy and the network. For each comparison, we show the associations as both scatterplots (top subpanels) and histograms (bottom subpanels). **2A:** Pairwise associations between biomarkers across cohorts. **2B:** Associations between biomarkers and connectivity to the entorhinal cortex. **2C:** From left to right, associations between the change in tau over time and tau, amyloid, connectivity-mediated tau spreading, and connectivity-mediated amyloid spreading. For the latter two comparisons, we employed the simple association proposed by the well-known network diffusion model: 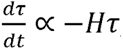, where H is the network Laplacian matrix^12^

### II. Evidence-based development and adjudication of competing hypotheses of protein aggregation

The above data snapshots provide empirical support to the hypothesis of a cross-species interaction between Aβ and tau; however, they do not reveal their exact mechanistic form, nor the mode of brain-wide protein propagation, whether connectome-mediated, proximity- or fiber distance-based. To explore various mechanistic hypotheses, we first developed a base model of network transmission of tau and Aβ (Eq (**1**)), where these two species evolve independently and migrate between regions via the connectome. We then extended this model to account for several different potential interactions and modes of transmission (**Table 1)**. See **SI:Note 1** for mathematical details.

### Group level empirical validation and model fitting

We next performed model fitting on empirical group data of each model initiated at the canonical EC-seeding of tau, using a robust maximum a posteriori (MAP) inference procedure we developed (detailed in **SI: Note 6)**. Cross-sectional group ADNI data were correlated against the fitted model’s evolution (**x_Aβ_**(*t*), ***x*_τ_**(*t*)) at every time *t*, and Pearson’s R was recorded. The resulting “R-t curves”, shown in **Figure 3** for the 1-way interaction *Aβ*→tau aggregation model, displayed a characteristic peak as more amyloid and tau pathology diffused into the network and increasingly recapitulated cross-sectional patterns. Subsequently model diverged from empirical pattern, decreasing R. Since our model posits that both amyloid and tau evolution happens on the same time axis, we report the model instant that maximized the posterior, rather than peak R for either tau or amyloid separately. The highest R value, *R_max_*, resulting from this “shared peak time”, called *t_max_*, is recorded for both amyloid and tau and considered as model evidence. Optimal fitted parameters shown in **Table S2** indicate substantial differences between groups. In total, we evaluated six theoretical interaction models (see **SI:Note 1** and **Table 1)**: **1)** No-interaction model (Eqs (**1**)); **2)** 1-way interaction model (Eqs (**2**)), tau affects amyloid aggregation but not vice versa; **3)** 1-way interaction, amyloid affects tau aggregation but not vice versa (Eqs **3**); **4)** Amyloid affects tau diffusion into the network, (Eqs (**4**)); **5)** 1-way (remote) interaction C · *Aβ*→tau aggregation, connectome-mediated *Aβ* induces tau aggregation at distant sites; and **6)** 2-way interaction model (Eq (**6**)). Each model was evaluated in identical fashion via MAP inference and identification of a single unique operating time *t_max_* that spans both amyloid and tau data. The numerical solutions of these models were evaluated on the canonical healthy connectome under the 86-region Desikan-Killiany parcellation. In order to assess model behavior broadly, in the following set of results we used a canonical model specification, given by the coarsely optimized (default) parameter values (refer to **SI: Note 6** and **Table S2** for details). Statistical comparison of these fitted models is shown in **Table 2**. All reported R values that are moderately to highly significant as denoted by * (p < 0.01) and ** (p < 0.001), **corrected for multiple comparisons.** The Akaike Information Criterion (AIC) is also presented to compare models of different complexity.

**Figure 3.**
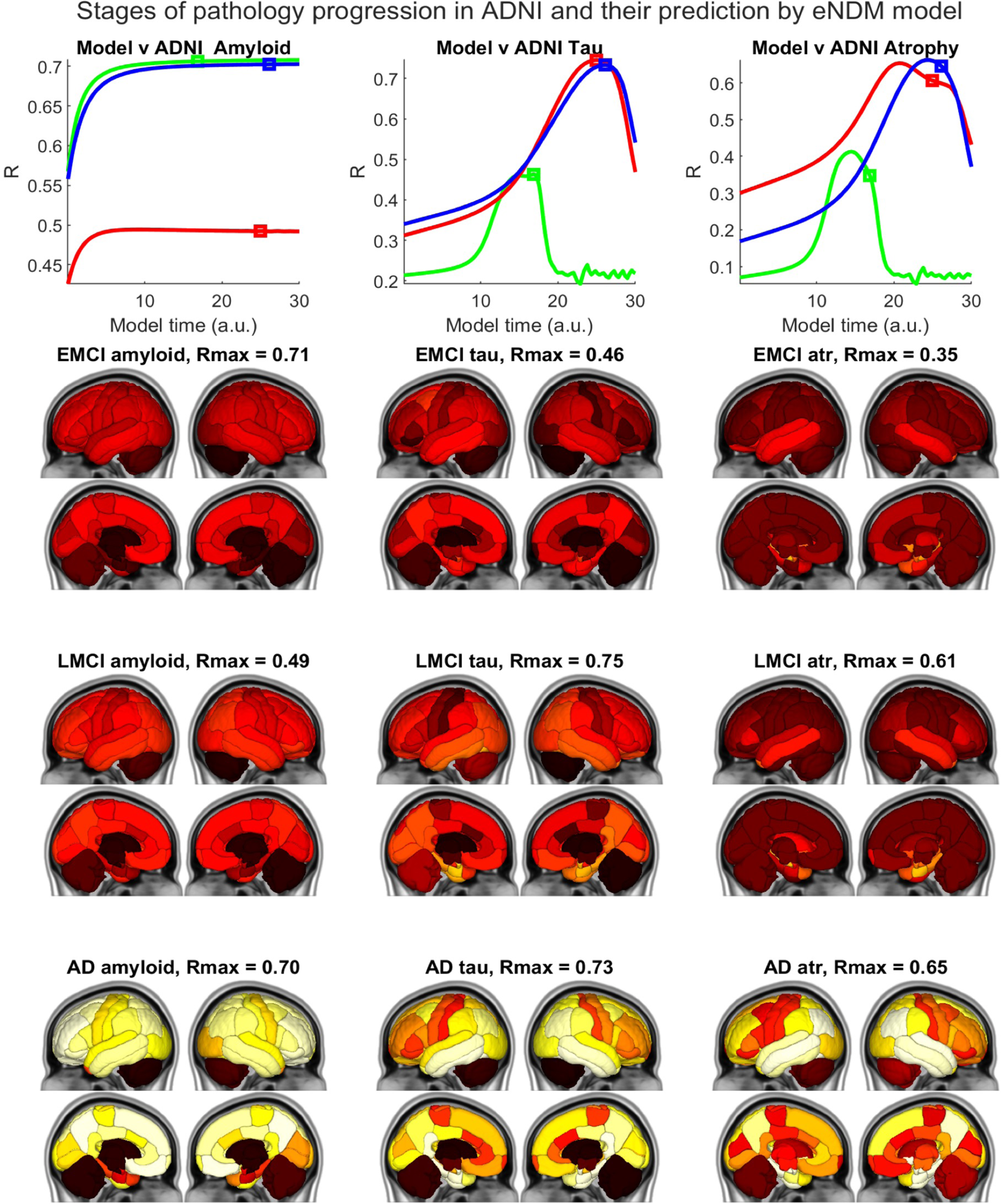
Validating evolution of model tau seeded at EC against empirical data. *Left column:* The top panel shows the behavior of model evidence R against model time *t* for amyloid. The model time t_max_ at which the model was evaluated against empirical data is denoted by square markers. The “glass brain” renderings show empirical amyloid-PET SUVr patterns from all 3 diagnostic groups. The R statistic between the model and empirical data are shown alongside. *Middle column:* The top panel shows the R between the model and empirical tau. The cross-sectional empirical tau-PET SUVr patterns from all 3 diagnostic groups are shown. *Right column:* Results for MRI-derived atrophy.

**Table 2:** Peak Pearson’s R of correlation between model and empirical regional statistics. Six theoretical interaction models’ evaluations are presented: the No-interaction model, without the interaction term (*γ* = 0); the 1-way interaction, tau→*Aβ* aggregation, whereby tau affects amyloid but not vice versa; the 1-way interaction, *Aβ*→tau diffusion model, whereby amyloid influences the rate of diffusion of tau but not vice versa; the 1-way interaction, *Aβ*→tau aggregation model, whereby amyloid affects tau but not vice versa; the 1-way interaction, C · *Aβ*→tau aggregation model, whereby distant amyloid affects tau but not vice versa; and the 2-way interaction Aβ←→tau aggregation model where both tau and amyloid affect each other via *γ*. Highly significant correlations: ** (p < 0.001), moderately significant: * (p < 0.01), all reported post Bonferroni correction. Aikeke information criterion (AIC) is reported for each model, in brackets. The highest R and lowest AIC values are highlighted in **boldface.**

| Empirical data | No- interaction<br>Model (1) | 1-way<br>interaction,<br>$\tau \rightarrow A\beta$<br>aggregation (2) | 1-way<br>interaction,<br>$A\beta \rightarrow \tau$<br>diffusion (3) | 1-way<br>interaction,<br>$A\beta \rightarrow \tau$<br>aggregation (4) | 1-way (remote)<br>interaction<br>$C \cdot A\beta \rightarrow \tau$<br>aggregation (5) | 2-way<br>interaction<br>$A\beta \leftrightarrow \tau$ (6) |
| --- | --- | --- | --- | --- | --- | --- |
| EMCI $A\beta$ | 0.71 ** (163) | 0.71 ** (163) | 0.71 ** (163) | 0.71 ** (163) | 0.71 ** (163) | 0.71 ** (166) |
| LMCI $A\beta$ | 0.49 * (223) | 0.49 * (223) | 0.49 * (223) | 0.49 * (223) | 0.49 * (223) | 0.45 * (229) |
| AD $A\beta$ | 0.70 ** (333) | 0.70 ** (333) | 0.70 ** (333) | 0.70 ** (333) | 0.70 ** (333) | 0.70 ** (335) |
| EMCI $\tau$ | 0.33 (200) | 0.38 (202) | 0.35 (201) | 0.48 * (190) | 0.48 * (192) | 0.36 (203) |
| LMCI $\tau$ | 0.60 ** (232) | 0.60 ** (234) | 0.49 * (247) | 0.75 ** (206) | 0.73 ** (211) | 0.70 ** (219) |
| AD $\tau$ | 0.57 ** (322) | 0.51 * (332) | 0.52 * (331) | 0.73 ** (296) | 0.71 ** (301) | 0.65 ** (315) |
| EMCI atrophy | 0.34 (171) | 0.32 (176) | 0.33 (175) | 0.35 (174) | 0.33 (178) | 0.37 (175) |
| LMCI atrophy | 0.60 ** (158) | 0.60 ** (162) | 0.56 * (171) | 0.62 ** (158) | 0.61 ** (159) | 0.48 * (184) |
| AD atrophy | 0.52 * (316) | 0.38 (331) | 0.39 (330) | 0.66 ** (299) | 0.62 ** (304) | 0.58 ** (312) |

#### Amyloid

Correlations between model and empirical AV45 SUVr are shown in **Table 2**. We found that all models worked equally well high significance for all three cohorts (*R_max_* = 0.71, *0.49*, and *0.70* for EMCI, LMCI, and AD, respectively), indicating that tau feedback onto amyloid is not required to explain its patterns of deposition at any stage of disease. This might reflect the well-known plateau effect of amyloid – also indicated by R-t curve (**Figure 3**, top left) that reaches a peak and then plateaus. Note, model time *t* has arbitrary units that may not directly correspond to empirical duration in years.

#### Tau

The network propagation of modeled tau starting from its seeding in EC was computed and its correspondence to regional group-average empirical AV1451-PET was assessed (**Table 2)**. In contrast to amyloid, we found significant differences between different interaction models. The 1-way interaction (*Aβ*→tau aggregation) was the most accurate and significant (post-Bonferroni) predictor of empirical tau-PET for all three cohorts: (EMCI: *R* = 0.51, p < 0.01; LMCI: *R* = 0.75, p < 0.001; AD: *R* = 0.73, p < 0.001), and achieved the lowest AIC. Only the 1-way (remote) interaction *Aβ*→tau model exhibited comparable correlation, but at the cost of additional model complexity hence lower AIC. None of the other proposed interaction models consistently outperformed the no interaction model (as assessed by AIC). Also unlike amyloid, the R-t curves of tau for 1-way interaction *Aβ*→tau aggregation model showed a slow and steady rise and a distinct late peak without a plateau effect (**Figure 3**, middle column). Correlation with EMCI group is poorer than other groups, likely due to lower PET uptake and inter-subject heterogeneity. These conclusions were further substantiated by performing pairwise t-tests between models following the Fisher’s R-to-z correction (**Table S3)**.

#### Atrophy

Since tau is highly co-localized with regional atrophy, we also show a comparison with MRI-derived group atrophy. However, we did not fit the model again to atrophy, instead borrowing the tau-fitted models for each cohort, under the assumption that tau is the primary, and atrophy is a surrogate for tau. Overall, the atrophy results were similar to but slightly weaker than tau-PET results. For the EMCI cohort, associations were similar for all models, indicating that the no interaction model was sufficient for explaining early neurodegeneration patterns (**Table 2)**; this is likely due to the low effect size measurable on MRI in this cohort. Associations with LMCI and AD were highly significant (*p* < 0.001) for the best model, 1-way interaction *Aβ*→tau, with *R* = 0.62 for LMCI and *R* = 0.66 for AD. However, LMCI atrophy was more or less equivalently associated with the no interaction and each of the three 1-way interaction Aβ→tau models. These conclusions were further substantiated by pairwise Fisher’s R-to-z t-tests between models (**Table S3)**.

#### Time of peak

The *t_max_* of maximum posterior followed the expected order: *t_max_* for AD > LMCI > EMCI (**Figure 3**, top). It is challenging to fit an accurate time axis to empirical data, since it does not have a measure of pathology duration – which may never be known. Interestingly, the peak for tau occurs more than a decade (in model “years”) after the plateau seen for amyloid – perhaps recapitulating well-known clinical and pathological examinations that suggest a decade-long delay between the two processes.

### Illustration of base model of Aβ and tau with no interactions

To get a qualitative picture, we generated surface renderings of the evolution of theoretical regional amyloid distribution under the no-interaction model as it propagates into the structural network (**Figure 4A**, left). Modeled amyloid evolution appeared to recapitulate the classic amyloid progression as proposed by Thal et al.^2^ and amyloid PET patterns^3^, proceeding from medial frontal and precuneus (areas with high baseline metabolism) into the wider network, only slowly entering temporal cortices. The middle column shows the evolution of tau on the same network under the no interaction model, starting from seeding event in the bilateral EC. This models largely stays within temporal cortices, with low but non-zero spread into other regions.

**Figure 4.**
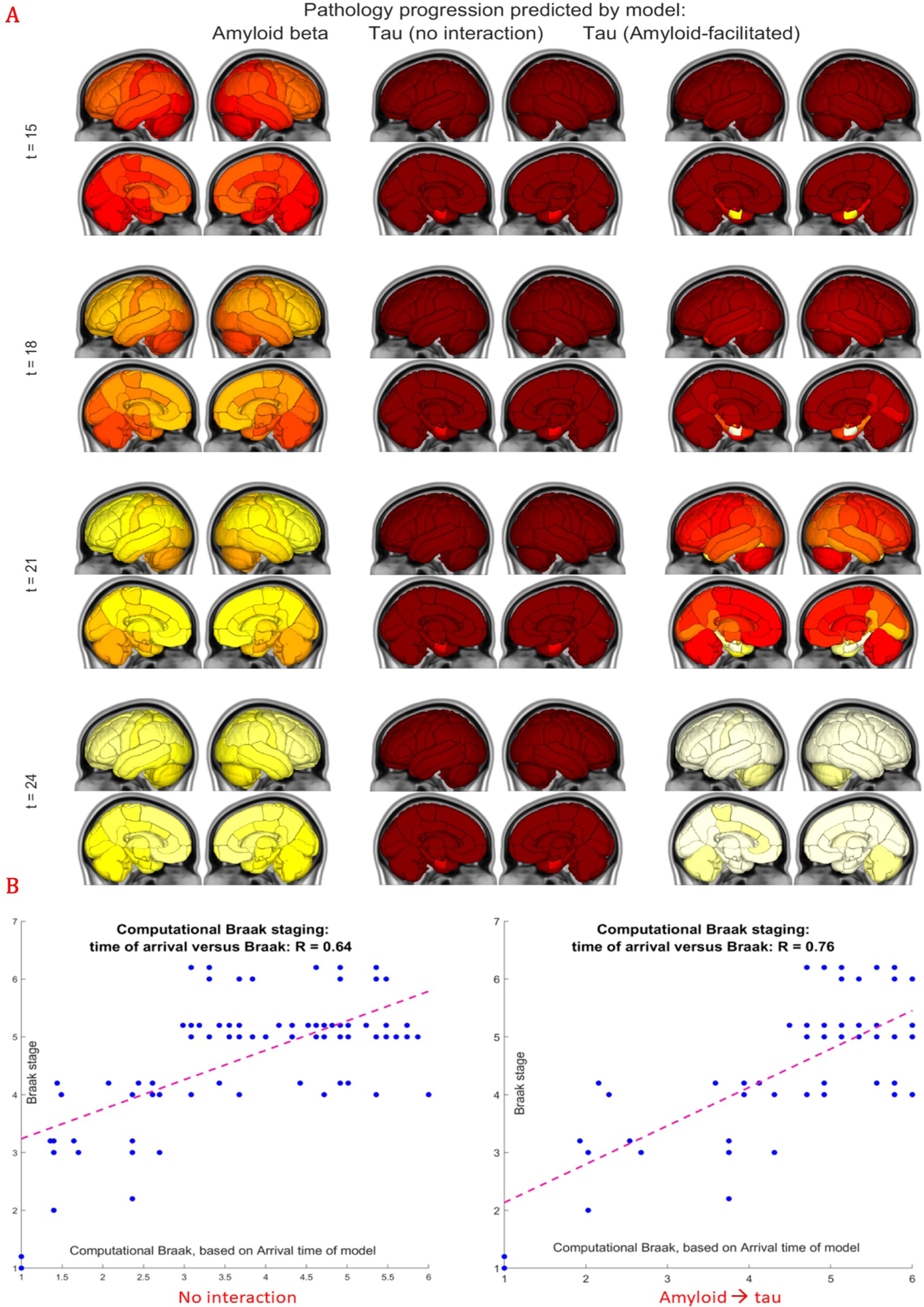
The evolution of model dynamics of amyloid and EC-seeded tau. **A**: Spatiotemporal evolution of the model evaluated by numerical integration of Eqs (1a, 2a), on the 86x86 healthy Desikan connectome using default model parameters. The first column shows the evolution of amyloid, the second column of tau only, and the third column of amyloid-facilitated tau. Each sphere represents a brain region, its diameter is proportional to pathology burden at the region, and is color coded by lobe (blue = frontal, purple = parietal, red = occipital, green = temporal, cyan = cingulate, black = subcortical). **B**: Prediction of Braak stages using computational model using a 6-stage “computational Braak” staging system based on the time-of-arrival calculation of each group’s fitted eNDM model, shown for the no interaction model (left) and 1-way interaction Aβ→tau aggregation model.

### Network transmission of Aβ and tau with cross-species interaction

Of the 6 interaction models encompassing various mechanistic hypotheses, here we detail the best interaction model (*Aβ*→tau aggregation)^16–18^, illustrated in **Figure 4A** right. Model tau remained confined to the medial temporal lobe until sufficient levels of Aβ had spread to and accumulated there. There onward, in contrast to the “pure tau” evolution, it took on a more aggressive trajectory, spreading first to nearby limbic, then basal forebrain, then parietal, lateral occipital and other neocortical areas, in close concordance with Braak’s six tau stages^1^ (**Figure 4B)**. Thus, the facilitation of tau by amyloid does not lead to colocalization of the two until late stages, helping explain why regional Aβ patterns do not coincide with tau and atrophy patterns. Conversely, the absence of the interaction term led modeled tau to remain confined to the temporal lobe (**Figure 4A**, middle), mirroring primary age-related tauopathy (PART), a new classification for mild neurofibrillary degeneration in the medial temporal lobe, but no Aβ plaques^55^.

**Figure S3** shows the global accumulation of theoretical pathology over model time, evaluated at default parameters. All proteins increase over time, but amyloid-facilitated tau diverges dramatically from the non-facilitated tau at around t=15, mirroring **Figure 4. Figure S3** makes it clear that while the “pure” tau model also captures empirical data, it does so far slower and achieves far less correlation strength than the facilitated version. Additionally, for comparison, the AAL-connectome-based model evolution is shown in **Figure S4,** with very similar behavior, indicating that the choice of atlas or processing pipeline did not drive results.

### Predicting Braak stages using computational model

Using the time-of-arrival calculation of each group’s fitted eNDM model, we developed a 6-stage “computational Braak” staging system. The predicted Braak stages strongly agree with the original Braak stages, applied to the DK atlas parcellation, achieving *R* = 0.76, p < 10^-6^ for the best-adjudicated model (**Figure 4B**, right). In comparison, the non-interacting model (**Figure 4B**, left), while being significant at *R* = 0.64, is substantially worse, suggesting that <u>amyloid→ tau interaction is a necessary factor in Braak staging</u>.

### Permutation testing to demonstrate disease specificity

Various permutation tests were deployed to demonstrate that the presented model only recapitulates empirical regional distributions when it is applied in the correct region order and to the correct human connectome, detailed in **SI: Note 7.** Under 500 random permutations of atrophy and tau (**Figure S5-A)**, and of the connectome itself (**Figure S5-B)** the above-reported R values remain highly significant compared to these “null” distributions (*p* < 10^-3^ for all groups). This was true whether the Pearson or Spearman correlation was used as the performance metric (**Figure S6)**.

### Comparison of different modes of spread

We implemented two alternative modes of spread (**Table 1)**: **1)** Pathology spread depends only on the shortest Euclidean distance; and **2)** transmission between regions is inversely proportional to the average length of fiber projections between regions. See **Methods** and quantitative comparison in **Table 3**. The model in each case was refitted using the MAP estimator as before, hence each fitted model may represent the best-case scenario for that hypothesis. We used paired t-tests following Fisher’s R-to-z transformation to statistically compare different spread models, while accounting for the correlation between dependent variables. We found that: **1)** For both tau and amyloid evolution, connectome-mediated spread model gave the closest correspondence to empirical data; and **2)** Euclidean spatial spread was slightly but not significantly superior to fiber distance-based spread. For these comparisons we chose the best interaction model identified in **Table 1** (1-way *Aβ* → tau aggregation) for all three networks. However, we repeated these comparisons using other interaction terms as well, and did not find significant differences in performance (Fisher’s p-value = N.S.).

**Table 3:** Pearson’s R of correlation between alternative spread models and empirical regional statistics. Fisher’s R-to-z transform was computed between the connectome and the alternative modes of spread, and its p-value is shown alongside. The connectome-mediated model significantly outperforms the two distance-based models (p values denoted in red font).

| Empirical data | *Aβ* → *tau*<br>interaction,<br>Connectivity | Fiber<br>Distance | Fisher R-to-z,<br>Fiber vs C | Euclidean<br>Distance | Fisher R-to-z,<br>Euclidean vs C |
| --- | --- | --- | --- | --- | --- |
| EMCI Aβ | 0.71 | 0.56 | **p < 0.01** | 0.57 | **p < 0.01** |
| LMCI Aβ | 0.49 | 0.43 | p = 0.10 | 0.42 | p = 0.09 |
| AD Aβ | 0.70 | 0.56 | **p < 0.01** | 0.57 | **p < 0.01** |
| EMCI tau | 0.48 | 0.25 | **p < 0.01** | 0.24 | **p < 0.01** |
| LMCI tau | 0.75 | 0.31 | **p < 0.01** | 0.35 | **p < 0.01** |
| AD tau | 0.73 | 0.34 | **p < 0.01** | 0.36 | **p < 0.01** |
| EMCI atrophy | 0.35 | 0.00 | **p < 0.01** | 0.18 | **p = 0.03** |
| LMCI atrophy | 0.62 | 0.29 | **p < 0.01** | 0.41 | **p = 0.025** |
| AD atrophy | 0.66 | 0.17 | **p < 0.01** | 0.22 | **p < 0.01** |

### Translational aspects

Having established the network model’s validity on group data, and having determined which modes of the tau-amyloid-network interactions are relevant and empirically supported, we showcase two key results related to translational aspects in patients: etiologic heterogeneity and the capacity to predict an individual subject’s spatiotemporal trajectory of AD.

#### Uncovering etiologic heterogeneity: Repeated seeding to assess alternative seeding sites

The entorhinal cortex (EC) was chosen above as the canonical tau seeding site. To establish other regions’ seeding plausibility, we repeatedly simulated the selected network interaction model seeded from every possible region bilaterally. For each seed region, peak Pearson’s R between model and ADNI tau-PET data is shown in **Figure 5A**. EC is among the best overall cortical seeding sites, while hippocampus (HP) is the best subcortical site. Other prominent seeding sites include parahippocampal gyrus (PHP) and fusiform gyrus (Fus), which are adjoining EC and appear frequently similar in tau uptake to EC on PET imaging. Thus the quantification of these structures as likely seeding locations affirms our model’s relevance and plausibility. The evolution of pathology from some of these non-EC sites is illustrated in **Figures S7** and **S8,** for HP-seeding and IT-seeding, respectively. While substantially similar to EC seeding of **Figure 3**, a notable difference is that HP seeding leads to higher involvement of medial temporal and subcortical structures. Of these four plausible seeding sites, EC is unique in giving consistently one of the highest seeding likelihood despite having lower levels of empirical tau deposition than others.

**Figure 5.**
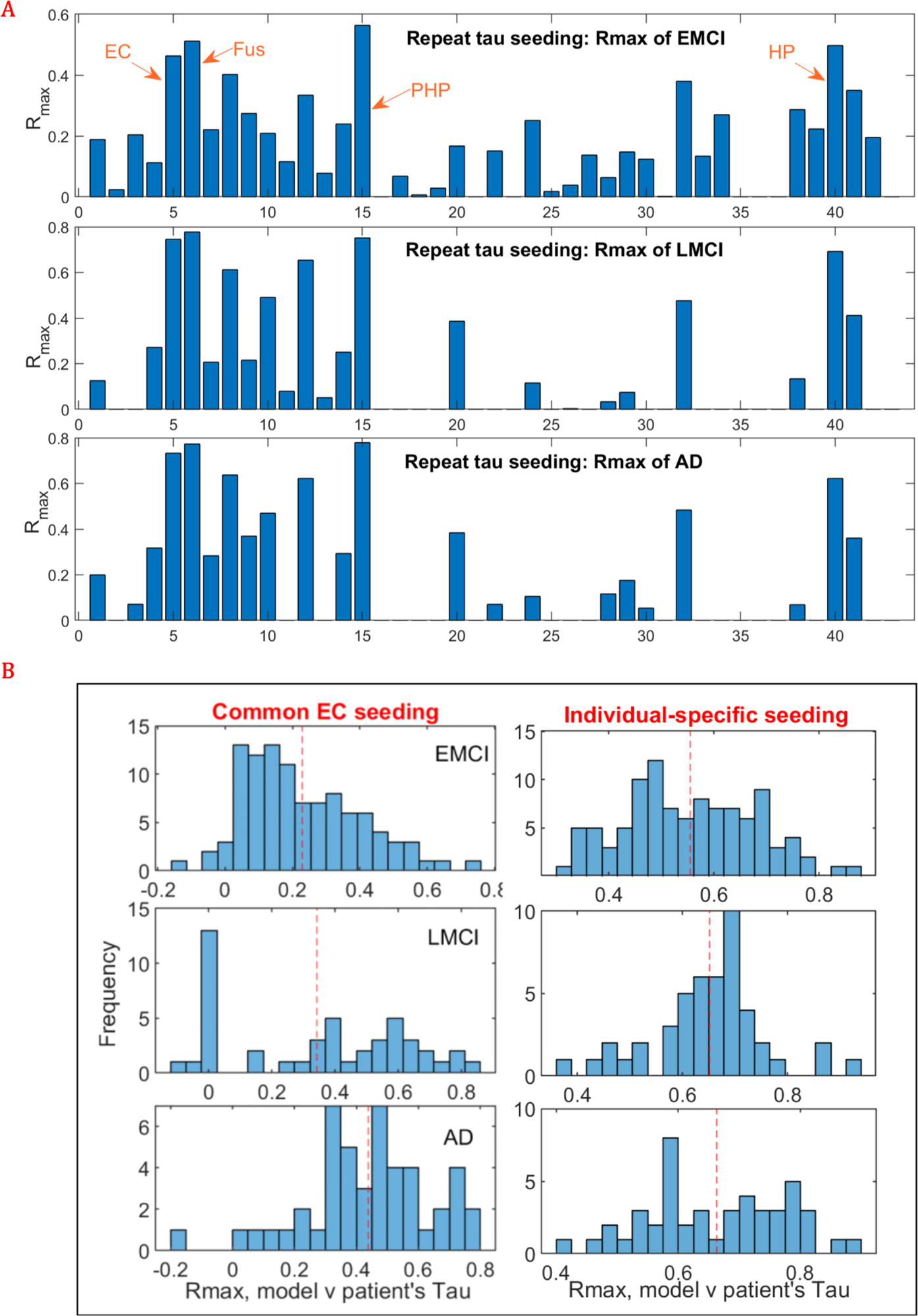
Translational applications. **A:** Bar charts of repeated seeding of each region in turn. For each seed region, peak Pearson’s R over model time, corresponding to the correlation between the best interaction model (amyloid-facilitated tau aggregation) and empirical tau PET data, is shown, for each of the 3 diagnostic groups: EMCI (top), LMCI (middle) and AD (bottom). Entorhinal cortex (EC) is amongst the best cortical seeding sites and Hippocampus (HP) the best subcortical site. Other likely seeding sites include Fusiform (Fus) and Parahippocampal gyrus (PHP). **B**: Histograms of *R_max_* achieved by fitting the model parameters to individual subjects’ baseline regional tau data, after imposing a common seeding site located at the EC (left) or after using the best seeding site for each subject (right). Here the best model combination from **Table 2** was used: 1-way interaction (Aβ → tau) on the structural connectivity graph, and the evaluation was performed on tau-PET scans in ADNI3. Average *R_max_* over all subjects within each diagnostic group is shown in red dashed line.

#### Individual subject fitting and prediction

For translational applications we must necessarily move away from group-wise to individual subjects, while accommodating both etiologic heterogeneity as well as potentially subject-specific model parametrization. In order to achieve this we deployed our robust inference procedure (see **SI: Note 6)** on individual subjects’ multimodal regional biomarkers from ADNI-3 (demographics in **Table S1)**, under the 1-way interaction *Aβ* → tau aggregation model. **Figure 5B** shows the histograms of *R_max_* between observed and model-predicted regional tau distribution in each subject. For comparison we show the results of both canonical (EC) seeding and each individual’s best seeding site. EC seeding was significantly worse (paired t-test after Fisher R-to-z: *p* < 10^-7^, corrected), implying that a common seeding site may not be appropriate for all subjects and revealing a potential etiological factor behind observed heterogeneity in AD-spectrum subjects. With individual-specific seeding and model fitting, we were able to achieve excellent prediction of the subjects’ tau distributions, with *R_max_* ranging widely up to a maximum of 0.9 and mean of 0.65 for LMCI and AD, and slightly lower for EMCI subjects. The mean *R_max_* across individuals in **Figure 5B** is similar to but slightly lower than the *R_max_* achieved on the group-averaged tau data of **Figure 4** and **Table 2**, which is expected due to heterogeneity as well as measurement noise in individual subjects. It is also noteworthy that canonical EC seeding fails in approximately half of the subjects in EMCI and LMCI cohorts (i.e. *R_max_* < 0.3) but only a small minority of diagnosed AD patients: indicating that higher stages have lower etiologic heterogeneity.

## DISCUSSION

The protein-protein interaction of amyloid and tau, commonly denoted by the A-T-N rubric^56^, is a central feature and key to understanding AD pathophysiology^17,19–21^, but has been difficult to reconcile with the observations of dissociated spatial distribution of tau and amyloid. The exact cause-effect mechanisms by which amyloid and tau regulate each other and cause downstream neurodegeneration and symptomatology remain poorly understood and empirically unsupported in human disease. Prior mechanistic studies in model organisms may not be entirely germane to human disease and empirical evidence for them is difficult if not impossible to acquire in AD patients. Yet, the interrogation of these aspects is critical to achieve disease understanding and future therapeutic options. In this study we took a computational rather than experimental approach to explore these issues directly in human AD. Our objectives were twofold: <u>First</u>, to test whether a mathematical encoding of network spread due to trans-neuronal proteopathic transmission^8,12,57^, combined with local interaction between amyloid and tau pathologies, is capable of recapitulating observed pathology progression in AD; <u>Second</u>, to infer quantitatively the factors of cross-species interactions, their causal direction, and their associated kinetic rate parameters in patients’ brains.

Our major findings are: <u>First</u>, the joint model successfully recapitulates the spatiotemporal progression of both amyloid and tau. The local production of Aβ driven by glucose metabolism followed by subsequent network spread correctly predicts the spatial distribution of empirical Aβ. Starting from EC, model tau predicts empirical tau, with prominence in temporal areas, followed by network ramification in limbic and wider cortices. <u>Second</u>, the best interaction model is one where Aβ influences tau aggregation, but not tau transmission. This one-way Aβ→tau interaction is an essential component of their spatiotemporal propagation predicted by the computational model, without which it does not fully recapitulate empirical amyloid and tau spatial patterns in patients. This model also excels in computationally predicting tau Braak stages. <u>Third</u>, 2-way interaction is worse than 1-way interaction in recapitulating empirical patterns, and the reverse interaction (tau→Aβ) does no better than the no interaction model. Using our “computational Braak” results, we showed that these alternate interactions are less consistent with Braak staging. These data constitute to our knowledge the first numerically rigorous evidence of the precise mode of amyloid-tau interaction in human AD. <u>Fourth</u>, connectome-mediated spread of tau outperforms other modes of spread, whether by proximity or by fiber length, statistically confirming hitherto descriptive observations that spatial or anisotropic diffusion are insufficient to correctly predict AD pathology. We verified these group-average results on individual subjects, both at baseline and longitudinally, and found essentially equivalent results.

The overall picture that emerges resembles the broad hypothesis posed at the beginning of the paper (**Figure 1A)**, but with a distinct inclination toward **amyloid-facilitation** rather than the classic amyloid-cascade hypothesis: Following diffuse production of amyloid in proportion to metabolism, and focal production of tau at the EC, aggregation into plaques and tangles occur at networked sites following graph topology. Tau pathology is further aggravated by amyloid, but not vice versa. Finally, classic, spatially divergent amyloid and tau patterns are established – frontal-dominant for amyloid and temporal dominant for tau. As discussed below, these results point to an underlying parsimony and universality, with the potential to explain several poorly understood aspects of AD progression.

### Network transmission drives divergent spatiotemporal progression of Aβ and tau

That the spatiotemporal patterning of Aβ, tau and atrophy are distinct is well-known but not fully understood^5^ and appears at odds with the prevailing amyloid cascade hypothesis^3,67^. Soluble Aβ is sometimes invoked to help explain the discrepancy, but this has been questioned^3^. Our first finding, that divergent evolution of theoretical amyloid and tau on the same network successfully recapitulates empirical progression, suggest that the governing mechanism behind these long-observed discrepancies may be related to network spread combined with metabolic drivers. Importantly, this spatial divergence holds even in the presence of amyloid-facilitation.

Indeed, we find that Aβ production driven by glucose metabolism and local APP pool, followed by a certain amount of network dissemination, significantly recapitulates regional amyloid deposition (**Figure 3)**: *R* = 0.71 (EMCI), *R* = 0.70 (AD). Neural activity is known to regulate the production and secretion of Aβ^3^. In transgenic mice, neural activity affects Aβ secretion^60^ through synaptic exocytosis^61^ and later deposition of plaques^58^, by modulating the release of cleavage products of APP^59^. In humans, Aβ release parallels fluctuations in synaptic activity in human sleep/wake cycles^62^. AV45 uptake is higher in hubs, multimodal cortices^63^ and default mode network, all characterized by higher baseline metabolism^64^. In contrast to Aβ, tau in our models proceeded outward from EC, to lateroinferior temporal cortices, thence to parietal, lateral occipital and medial frontal areas, in close concordance with Braak’s six tau stages^65^ (**Figure 4B)**. The best model yielded highly significant correlations against AV1451-PET data (**Figure 3)**: *R* = 0.75 (LMCI), *R* = 0.73 (AD). We found that the EC was the consensus best seeding site at both a group and individual level (**Figure 5)**, although substantial heterogeneity was apparent within each cohort. Further, the model correlated with MRI-derived regional atrophy, an excellent surrogate for underlying tau^43,66^, with high significance. The presented correlations are significantly stronger than 500 random permutations (**Figure S5)**, suggesting that the model is specific to the human connectome and brain topography.

### Aβ-facilitated tau aggregation plays a critical role in progression

Our second major result is that Aβ-tau interaction (see e.g.^47,48^) played an important role in forcing the extra-temporal spread of tau. Aβ influences the course and severity of tau and atrophy^16–18^. In our mathematical exposition, modeled tau seeded at EC remained confined to the temporal lobe until sufficient levels of Aβ had spread to and accumulated in temporal cortices (**Figure 4)**. The amyloid-facilitated model tau then started diverging from pure tau, taking on a more aggressive trajectory, spreading first to nearby limbic, then basal forebrain, then other neocortical areas. Thus, *the facilitation of tau by amyloid does not lead to colocalization of the two until late stages,* further helping to explain why regional Aβ patterns do not coincide with tau and atrophy patterns^76,77^. Conversely, the absence of the interaction term led modeled tau to remain confined to the temporal lobe (**Figure 4)**, mirroring primary age-related tauopathy (PART), a new classification for mild neurofibrillary degeneration in the medial temporal lobe, but no Aβ plaques ^55^. Thus, medial temporal NFTs may be involved in two divergent processes: AD and PART. Purely tau-specific abnormalities (e.g., the MAPT gene H1 haplotype) would predispose the subject to PART, whereas additional Aβ abnormalities (e.g., the dysregulation of presenilin, APP or APOE ε4 allele) would cause AD predisposition^55^. In the present work, the non-facilitated tau model evolved less rapidly and was less successful than amyloid-facilitated tau model (**Table 2**, **Figure 3)**. These results provide model-based support to prior neuroimaging observations: Sepulcre *et al* found several convergence zones in temporal and entorhinal cortex where amyloid and tau might interact^78,79^. Franzmeier *et al* showed that tau uptake in Aβ-controls was restricted to IT, but was observed in extra-temporal areas in Aβ+ subjects^80^. Remarkably, our theoretical modeling also gives the same regions (IT and EC – see **Figure 4)** as key areas of convergence between amyloid and tau, where the “arrival” of the former is accompanied by prominent deposition of the latter.

Ittner and Götz^17^ have suggested three modes of interaction: (1) Aβ drives tau pathology; (2) synergistic toxic effects of Aβ and tau; and (3) tau mediates Aβ toxicity. Aβ and pathological tau co-localize in synapses^68,69^, and tau is essential for Aβ-induced neurotoxicity^70^. Aβ seeded exogenously in tau transgenic mice elicited aggressive tau pathology in retrogradely connected regions^18^. Amyloid pathology accelerated tau deposition in double transgenic mice, but the reverse effect was not observed^16^. A “seminal cell biological event” in AD pathogenesis was suggested, whereby acute, tau-dependent loss of microtubule integrity is caused by exposure of neurons to readily diffusible Aβ^21^. However, the question remains controversial and other studies have suggested that tau propagation across connected regions is unlikely to be a gain of function mediated by the presence of Aβ^71^. Nonetheless, there are other mechanistic routes to enact this interaction, such as Aβ-related microglial activation promoting local tau-hyperphosphorylation^72^, enhancing tau spread across connected neurons^73^, PET imaging confirms early microglial activation in inferior temporal sites of tau accumulation^74^. Hence, an indirect mediation of Aβ→tau may occur via microglial activation; see review^75^.

### Alternative models of spread and interaction are less plausible

None of the other five theoretical interaction models (**Table 1)** compared favorably to *Aβ*→tau. This included amyloid effect on tau diffusion; and its (remote) interaction with tau aggregation (C · *Aβ*→tau) – two popular hypotheses in recent literature: We find little support for the role of amyloid in exacerbating tau spread, potentially mediated by microglia^47,49^. Remote effect of amyloid on tau was posited as a potential explanation of the dissociation between the two^5,40,50^. Sepulcre *et al* find that tau accumulation in temporal areas relates to “massive Aβ elsewhere in the brain”, linking both pathologies at the large-scale level^78,79^. However our analysis does not support this view. Surprisingly, the 2-way model (e.g.^51^) was insignificantly different from the 1-way model for both amyloid and tau, while they are identical for amyloid (**Table 2)**. The tau→amyloid interaction (e.g.^29^) performed relatively worse on tau data as well, suggesting that tau driving amyloid is not a clinically relevant factor in pathology ramification, and its addition gives poorer AIC scores. That amyloid results are identical for both modes was somewhat surprising, but this may be due to amyloid evolution peaking before a significant tau-influence was observed. Hence, by the time tau can begin to influence amyloid, the pear R against empirical amyloid distribution has already been reached.

Intriguingly, alternative spread models based on proximity^50,52,53^ or fiber distance^34,54^ did not give good correspondence with empirical tau, and were essentially identical for amyloid (**Table 3)**. The amyloid result mirrors our previous exploration in mouse data^81^, where too connectivity was not better than spatial spread. Taken together, our findings serve to rule out or deprioritize many alternative hypotheses and mechanisms of AD progression.

### Broader Implications and future work

The most important implication is that the best-adjudicated model (*Aβ*→tau interaction accompanied by network spread) gives a clear mechanistic target for future drug design. Scientifically, this study bolsters the hypothesis of trans-neuronal transmission by demonstrating its role in humans, an aspect that cannot be studied directly. Clinically, the new model could provide a unique opportunity for computational tracking and prediction of individual patients, especially after integrating multimodal imaging biomarkers (e.g., MRI, AV1451- and AV45-PET); we have provided proof-of-concept in **Figure 5**. Subject-specific seeding sites gave much higher accuracy than uniform common seeding of EC – revealing important inter-subject heterogeneity of etiology and ramification. Future model extensions are planned to incorporate machine learning and data-driven multifactorial^36^ and event-based models^37,38^, which give disease duration, not available here. Applications of the model as outcome measures in clinical drug trials are planned. Since trans-neuronal spread is a common feature of neurodegeneration, the best-adjudicated model may be equally applicable to co-morbid pathologies seen in other disorders like Parkinson’s, Lewy Body, frontotemporal and other dementias.

### Limitations

Neuroimaging software pipelines have several limitations in image resolution, noise and artifacts ^12^. DTI suffers from susceptibility artifacts and poor resolution. PET has poor resolution compared to MRI, and AV45 and AV1451 tracers show significant non-specific binding. Tractography can under-estimate crossing fibers and long tracts. Small subcortical structures can present challenges in inferring connectivity. This study was cross sectional hence longitudinal information was not utilized.

## ACKNOWLEDGEMENTS

Authors wish to acknowledge assistance in gathering ADNI data by Dr Duygu Tosun, Daren Ma and Areez Malik. This research was supported by the following grants from the National Institutes of Health: R01NS092802, R01EB022717, RF1AG062196, R56AG064873.

## COMPETING INTERESTS

The authors declare no conflict of interest.

## Notes

### Competing Interest Statement

The authors have declared no competing interest.

### Summary of Updates

Updated manuscript and supplement, with separate figure files

## REFERENCES

1. Braak, H. & Braak, E. Neuropathological stageing of Alzheimer-related changes. Acta Neuropathol. 82, 239– 259 (1991).

2. Thal, D. R., Rüb, U., Orantes, M. & Braak, H. Phases of A beta-deposition in the human brain and its relevance for the development of AD. Neurology 58, 1791–800 (2002).

3. Jagust, W. J. & Mormino, E. C. Lifespan brain activity, β-amyloid, and Alzheimer’s disease. Trends Cogn. Sci. 15, 520–6 (2011).

4. Hardy, J. & Selkoe, D. J. The amyloid hypothesis of Alzheimer’s disease: progress and problems on the road to therapeutics. Science 297, 353–6 (2002).

5. La Joie, R. et al. Region-specific hierarchy between atrophy, hypometabolism, and β-amyloid (Aβ) load in Alzheimer’s disease dementia. J. Neurosci. 32, 16265–73 (2012).

6. Rabinovici, G. D. et al. Increased metabolic vulnerability in early-onset Alzheimer’s disease is not related to amyloid burden. Brain 133, 512–28 (2010).

7. Jack Jr, C. R., et al. Hypothetical model of dynamic biomarkers of the Alzheimer’s pathological cascade. Lancet Neurol. 9, 119–128 (2010).

8. Jucker, M. & Walker, L. C. Self-propagation of pathogenic protein aggregates in neurodegenerative diseases. Nature 501, 45–51 (2013).

9. Frost, B. & Diamond, M. I. Prion-like mechanisms in neurodegenerative diseases. Nat. Rev. Neurosci. 11, 155– 9 (2010).

10. Clavaguera, F. et al. Transmission and spreading of tauopathy in transgenic mouse brain. Nat. Cell Biol. 11, 909–13 (2009).

11. Iba, M. et al. Tau pathology spread in PS19 tau transgenic mice following locus coeruleus (LC) injections of synthetic tau fibrils is determined by the LC’s afferent and efferent connections. Acta Neuropathol. 130, 349– 62 (2015).

12. Raj, A., Kuceyeski, A. & Weiner, M. A network diffusion model of disease progression in dementia. Neuron 73, 1204–15 (2012).

13. Zhou, J., Gennatas, E. D., Kramer, J. H., Miller, B. L. & Seeley, W. W. Predicting regional neurodegeneration from the healthy brain functional connectome. Neuron 73, 1216–27 (2012).

14. Pievani, M., de Haan, W., Wu, T., Seeley, W. W. & Frisoni, G. B. Functional network disruption in the degenerative dementias. Lancet. Neurol. 10, 829–43 (2011).

15. Raj, A. et al. Network Diffusion Model of Progression Predicts Longitudinal Patterns of Atrophy and Metabolism in Alzheimer’s Disease. Cell Rep. **in print**, 359–369 (2015).

16. Hurtado, D. E. et al. A{beta} accelerates the spatiotemporal progression of tau pathology and augments tau amyloidosis in an Alzheimer mouse model. Am. J. Pathol. 177, 1977–88 (2010).

17. Ittner, L. M. & Götz, J. Amyloid-β and tau--a toxic pas de deux in Alzheimer’s disease. Nat. Rev. Neurosci. 12, 65–72 (2011).

18. Götz, J., Chen, F., van Dorpe, J. & Nitsch, R. M. Formation of neurofibrillary tangles in P301l tau transgenic mice induced by Abeta 42 fibrils. Science 293, 1491–5 (2001).

19. Walker, L. C., Lynn, D. G. & Chernoff, Y. O. A standard model of Alzheimer’s disease? Prion vol. 12 261–265 (2018).

20. Kara, E., Marks, J. D. & Aguzzi, A. Toxic Protein Spread in Neurodegeneration: Reality versus Fantasy. Trends in Molecular Medicine vol. 24 1007–1020 (2018).

21. King, M. E. et al. Tau-dependent microtubule disassembly initiated by prefibrillar β-amyloid. J. Cell Biol. (2006) doi:10.1083/jcb.200605187.

22. Spires-Jones, T. L. & Hyman, B. T. The Intersection of Amyloid Beta and Tau at Synapses in Alzheimer’s Disease. Neuron 82, 756–771 (2014).

23. He, Z. et al. Amyloid-β plaques enhance Alzheimer’s brain tau-seeded pathologies by facilitating neuritic plaque tau aggregation. Nat. Med.24, 29–38 (2018).

24. Maphis, N. et al. Reactive microglia drive tau pathology and contribute to the spreading of pathological tau in the brain. Brain 138, 1738–1755 (2015).

25. Mancuso, R. et al. CSF1R inhibitor JNJ-40346527 attenuates microglial proliferation and neurodegeneration in P301S mice. Brain 142, 3243–3264 (2019).

26. Asai, H. et al. Depletion of microglia and inhibition of exosome synthesis halt tau propagation. Nat. Neurosci. 18, 1584–1593 (2015).

27. Perez-Nievas, B. G. et al. Dissecting phenotypic traits linked to human resilience to Alzheimer’s pathology. Brain 136, 2510–2526 (2013).

28. Wu, J. W. et al. Neuronal activity enhances tau propagation and tau pathology in vivo. Nat. Neurosci. 19, 1085–1092 (2016).

29. Jackson, R. J. et al. Human tau increases amyloid β plaque size but not amyloid β-mediated synapse loss in a novel mouse model of Alzheimer’s disease. Eur. J. Neurosci. 44, 3056–3066 (2016).

30. Pooler, A. M., Phillips, E. C., Lau, D. H. W., Noble, W. & Hanger, D. P. Physiological release of endogenous tau is stimulated by neuronal activity. EMBO Rep. 14, 389–394 (2013).

31. Raj, A., Kuceyeski, A. & Weiner, M. A Network Diffusion Model of Disease Progression in Dementia. Neuron 73, 1204–1215 (2012).

32. Zhou, J., Gennatas, E. D., Kramer, J. H., Miller, B. L. & Seeley, W. W. Predicting regional neurodegeneration from the healthy brain functional connectome. Neuron 73, 1216–27 (2012).

33. Weickenmeier, J., Jucker, M., Goriely, A. & Kuhl, E. A physics-based model explains the prion-like features of neurodegeneration in Alzheimer’s disease, Parkinson’s disease, and amyotrophic lateral sclerosis. J. Mech. Phys. Solids 124, 264–281 (2019).

34. Fornari, S., Schäfer, A., Jucker, M., Goriely, A. & Kuhl, E. Prion-like spreading of Alzheimer’s disease within the brain’s connectome. J. R. Soc. Interface 16, (2019).

35. Iturria-Medina, Y., Sotero, R. C., Toussaint, P. J. & Evans, A. C. Epidemic Spreading Model to Characterize Misfolded Proteins Propagation in Aging and Associated Neurodegenerative Disorders. PLoS Comput. Biol. 10, e1003956 (2014).

36. Iturria-Medina, Y., Carbonell, F. M., Sotero, R. C., Chouinard-Decorte, F. & Evans, A. C. Multifactorial causal model of brain (dis)organization and therapeutic intervention: Application to Alzheimer’s disease. Neuroimage 152, 60–77 (2017).

37. Young, A. L. et al. A data-driven model of biomarker changes in sporadic Alzheimer’s disease. Brain 137, 2564–2577 (2014).

38. Oxtoby, N. P. et al. Data-Driven Sequence of Changes to Anatomical Brain Connectivity in Sporadic Alzheimer’s Disease. Front. Neurol. 8, 580 (2017).

39. Behrens, T. E. J., Berg, H. J., Jbabdi, S., Rushworth, M. F. S. & Woolrich, M. W. Probabilistic diffusion tractography with multiple fibre orientations: What can we gain? Neuroimage 34, 144–55 (2007).

40. Pandya, S., Kuceyeski, A., Raj, A. & Alzheimer’s Disease Neuroimaging Initiative. The Brain’s Structural Connectome Mediates the Relationship between Regional Neuroimaging Biomarkers in Alzheimer’s Disease. J. Alzheimer’s Dis. 55, 1639–1657 (2016).

41. Weiner, M. W. et al. The Alzheimer’s Disease Neuroimaging Initiative: a review of papers published since its inception. Alzheimers. Dement. 8, S1–68 (2012).

42. Glasser, M. F. et al. The minimal preprocessing pipelines for the Human Connectome Project. Neuroimage 80, 105–24 (2013).

43. Whitwell, J. L. et al. MRI correlates of neurofibrillary tangle pathology at autopsy: a voxel-based morphometry study. Neurology 71, 743–9 (2008).

44. Iba, M. et al. Synthetic tau fibrils mediate transmission of neurofibrillary tangles in a transgenic mouse model of Alzheimer’s-like tauopathy. J. Neurosci. 33, 1024–37 (2013).

45. Boluda, S. et al. Differential induction and spread of tau pathology in young PS19 tau transgenic mice following intracerebral injections of pathological tau from Alzheimer’s disease or corticobasal degeneration brains. Acta Neuropathol. 129, 221–37 (2015).

46. Kaufman, S. K. et al. Tau Prion Strains Dictate Patterns of Cell Pathology, Progression Rate, and Regional Vulnerability In Vivo. Neuron 92, 796–812 (2016).

47. He, Z. et al. Amyloid-β plaques enhance Alzheimer’s brain tau-seeded pathologies by facilitating neuritic plaque tau aggregation. Nat. Med.24, 29–38 (2018).

48. Gratuze, M. et al. Impact of TREM2R47H variant on tau pathology–induced gliosis and neurodegeneration. J. Clin. Invest. 130, 4954–4968 (2020).

49. Bolmont, T. et al. Induction of tau pathology by intracerebral infusion of amyloid-beta-containing brain extract and by amyloid-beta deposition in APP x Tau transgenic mice. Am. J. Pathol. 171, 2012–2020 (2007).

50. Sepulcre, J., et al. In Vivo Tau, Amyloid, and Gray Matter Profiles in the Aging Brain. J. Neurosci. 36, 7364–7374 (2016).

51. Pooler, A. M. et al. Tau - amyloid interactions in the rTgTauEC model of early Alzheimer’s disease suggest amyloid induced disruption of axonal projections and exacerbated axonal pathology. J. Comp. Neurol. 521, (2013).

52. Mezias, C. & Raj, A. Analysis of Amyloid-β pathology spread in mouse models suggests spread is driven by spatial proximity, not connectivity. Front. Neurol. 8, (2017).

53. Sepulcre, J., Sabuncu, M. R., Becker, A., Sperling, R. & Johnson, K. A. In vivo characterization of the early states of the amyloid-beta network. Brain 136, 2239–52 (2013).

54. Schäfer, A., Mormino, E. C. & Kuhl, E. Network Diffusion Modeling Explains Longitudinal Tau PET Data. Front. Neurosci. 14, (2020).

55. Crary, J. F. et al. Primary age-related tauopathy (PART): a common pathology associated with human aging. Acta Neuropathol. 128, 755–66 (2014).

56. Jack, C. R. et al. A/T/N: An unbiased descriptive classification scheme for Alzheimer disease biomarkers. Neurology (2016) doi:10.1212/WNL.0000000000002923.

57. Seeley, W. W., Crawford, R. K., Zhou, J., Miller, B. L. & Greicius, M. D. Neurodegenerative diseases target large-scale human brain networks. Neuron 62, 42–52 (2009).

58. Bero, A. W. et al. Neuronal activity regulates the regional vulnerability to amyloid-β deposition. Nat. Neurosci. 14, 750–756 (2011).

59. Nitsch, R. M., Farber, S. A., Growdon, J. H. & Wurtman, R. J. Release of amyloid beta-protein precursor derivatives by electrical depolarization of rat hippocampal slices. Proc. Natl. Acad. Sci. U. S. A.90, 5191–3 (1993).

60. Kamenetz, F. et al. APP processing and synaptic function. Neuron 37, 925–37 (2003).

61. Cirrito, J. R. et al. Synaptic activity regulates interstitial fluid amyloid-beta levels in vivo. Neuron 48, 913–22 (2005).

62. Gilestro, G. F., Tononi, G. & Cirelli, C. Widespread Changes in Synaptic Markers as a Function of Sleep and Wakefulness in Drosophila. Science (80-.). 324, 109–112 (2009).

63. Buckner, R. L. et al. Cortical hubs revealed by intrinsic functional connectivity: mapping, assessment of stability, and relation to Alzheimer’s disease. J. Neurosci. 29, 1860–73 (2009).

64. Buckner, R. L. et al. Molecular, structural, and functional characterization of Alzheimer’s disease: evidence for a relationship between default activity, amyloid, and memory. J. Neurosci. 25, 7709–17 (2005).

65. Braak, H. & Braak, E. Evolution of the neuropathology of Alzheimer’s disease. Acta Neurol. Scand. Suppl. 165, 3–12 (1996).

66. Vemuri, P. et al. Antemortem MRI based STructural Abnormality iNDex (STAND)-scores correlate with postmortem Braak neurofibrillary tangle stage. Neuroimage 42, 559–67 (2008).

67. Fjell, A. M. & Walhovd, K. B. Neuroimaging results impose new views on Alzheimer’s disease--the role of amyloid revised. Mol. Neurobiol. 45, 153–72 (2012).

68. Fein, J. A. et al. Co-localization of amyloid beta and tau pathology in Alzheimer’s disease synaptosomes. Am. J. Pathol. 172, 1683–92 (2008).

69. Takahashi, R. H., Capetillo-Zarate, E., Lin, M. T., Milner, T. A. & Gouras, G. K. Co-occurrence of Alzheimer’s disease ß-amyloid and τ pathologies at synapses. Neurobiol. Aging 31, 1145–52 (2010).

70. Amadoro, G. et al. Endogenous Aβ causes cell death via early tau hyperphosphorylation. Neurobiol. Aging 32, 969–90 (2011).

71. Franzmeier, N. et al. Functional brain architecture is associated with the rate of tau accumulation in Alzheimer’s disease. Nat. Commun. 11, 1–17 (2020).

72. Heurtaux, T. et al. Microglial activation depends on beta-amyloid conformation: role of the formylpeptide receptor 2. J. Neurochem. 114, 576–586 (2010).

73. Maphis, N. et al. Reactive microglia drive tau pathology and contribute to the spreading of pathological tau in the brain. Brain 138, 1738–1755 (2015).

74. Hamelin, L. et al. Early and protective microglial activation in Alzheimer’s disease: a prospective study using 18 F-DPA-714 PET imaging. Brain 139, 1252–1264 (2016).

75. Busche, M. A. et al. Tau impairs neural circuits, dominating amyloid-β effects, in Alzheimer models in vivo. Nat. Neurosci. 22, 57–64 (2019).

76. Vemuri, P. et al. MRI and CSF biomarkers in normal, MCI, and AD subjects: predicting future clinical change. Neurology 73, 294–301 (2009).

77. Jack, C. R. & Holtzman, D. M. Biomarker Modeling of Alzheimer’s Disease. Neuron 80, 1347–1358 (2013).

78. Sepulcre, J. et al. In vivo tau, amyloid, and gray matter profiles in the aging brain. J. Neurosci. 36, 7364–7374 (2016).

79. Marquié, M. et al. Validating novel tau positron emission tomography tracer [F-18]-AV-1451 (T807) on postmortem brain tissue. Ann. Neurol. 78, 787–800 (2015).

80. Franzmeier, N. et al. Functional connectivity associated with tau levels in ageing, Alzheimer’s, and small vessel disease. Brain 142, 1093–1107 (2019).

81. Mezias, C. & Raj, A. Analysis of Amyloid-β Pathology Spread in Mouse Models Suggests Spread Is Driven by Spatial Proximity, Not Connectivity. Front. Neurol. 8, 653 (2017).

82. Kuceyeski, A., Maruta, J., Relkin, N. & Raj, A. The Network Modification (NeMo) Tool: elucidating the effect of white matter integrity changes on cortical and subcortical structural connectivity. Brain Connect. 3, 451–63 (2013).

83. Tzourio-Mazoyer, N. et al. Automated anatomical labeling of activations in SPM using a macroscopic anatomical parcellation of the MNI MRI single-subject brain. Neuroimage 15, 273–89 (2002).

84. Alemán-Gómez, Y., Melie-García, L. & Valdés-Hernandez, P. IBASPM: Toolbox for automatic parcellation of brain structures. in Presented at the 12th Annual Meeting of the Organization for Human Brain Mapping (2005).

85. Shinohara, M. et al. Regional distribution of synaptic markers and APP correlate with distinct clinicopathological features in sporadic and familial Alzheimer’s disease. Brain 137, 1533–49 (2014).

86. Trabzuni, D. et al. MAPT expression and splicing is differentially regulated by brain region: relation to genotype and implication for tauopathies. Hum. Mol. Genet.21, 4094–103 (2012).

87. Villemagne, V. L. et al. Amyloid β deposition, neurodegeneration, and cognitive decline in sporadic Alzheimer’s disease: a prospective cohort study. Lancet Neurol. 12, 357–67 (2013).

