## Supplemental Material for "Understanding the complex interplay between tau, amyloid and the network in the spatiotemporal progression of Alzheimer’s Disease"

**This section includes:**

Supplemental Figures (**Figures S1** to **S8**)

Supplemental **Tables S1, S2, S3**

Supplemental Experimental Procedures (**Notes 1,2,3,6**)

Extended Data and statistics (**Note 4,5, 7-9**)

The full mathematical network-interaction model, and all its variants tested here, are given in **SI: Note 1**. Other pertinent methodological details and image processing pipelines are described in **Notes 2,3**, while Bayesian inference procedures are described in Note 6. Extended data and statistics are contained in the remaining **Notes**.

Supplemental Methods, Tables and Figures

**Note 1**. **Mathematical Model of Network Transmission**

**Notation**. Brain’s anatomic connectivity network is defined on the graph $\mathcal{G=\{V,E\}}$ whose nodes $v_{i}\mathcal{\in V,}i\in[1,N_{roi}]$ represent grey matter structures, and edges $e_{i,j}\mathcal{\in E}$ represent fiber connectivity. Structures $v_{i}$ comes from parcellation of brain MRI with $N_{roi}$ gray matter regions, and connection strength of the edge between any pair of regions i,j, $c_{i,j}$ is measured by fiber tractography ^39^, giving the network connectivity matrix $C=\{c_{i,j}, (i,j\mathcal{)\in E\}}$. Each node has an in-degree and an out-degree given by row and column sums of the matrix C: $\mathbf{d}_{row,i}= \sum_{j} c_{i,j}, \mathbf{d}_{col,j}= \sum_{i} c_{i,j}$. We also define the diagonal elements $c_{i,i}$to be 0.

Define $H$ as the graph Laplacian matrix $H=I-diag\left( \mathbf{d}_{row}\cdot\mathbf{d}_{col} \right)^{-1/2} C$. This Laplacian definition normalizes each node’s connectivity by the geometric mean of its in/out-degree, to accommodate brain parcellations of varying size and connectivity, and emphasizes nodes with high imbalance between incoming and outgoing connections. We define two time-varying vectors containing levels of $A\beta$ and tau at each node: $\mathbf{x}_{A\beta}(t)$ and $\mathbf{x}_{\tau}(t)$ respectively.

**Non-linear reaction + network diffusion with local growth and cross-tau-Aβ interaction**

Here we propose a complete and realistic model of pathology dynamics that incorporates the production, growth and spread of both Aβ and tau, as well as the interaction between them.

1) **The No-interaction model**:

Evolution of tau:	$\frac{d\mathbf{x}_{\boldsymbol{\tau}}(t)}{dt}= -\beta H\mathbf{x}_{\boldsymbol{\tau}}\left( t \right)+ {\alpha f}_{\tau}\left( t \right)\mathbf{e}_{ERC}$	(**1a**)

Evolution of Aβ:	$\frac{d\mathbf{x}_{\boldsymbol{A\beta}}(t)}{dt}= -\beta H\mathbf{x}_{\boldsymbol{A\beta}}\left( t \right)+ {\alpha f}_{A\beta}\left( t \right)(\mathbf{m}\cdot\mathbf{x}_{APP})$	(**1b**)

The first term on the right hand sides of Eqs (**1**) above represents network diffusion, following our prior work ^12^, whereby pathology spread follows regional concentration gradients restricted along network connections. This involves the connectome’s Laplacian matrix H and the diffusivity rate constant $\beta$, thus implying that its rate depends both on concentration gradients and the amount of connectivity between regions. This model captures trans-neuronal propagation as a connectivity- rather than distance-based process, as previous experimental work has suggested (Clavaguera 2009). Fiber length does not enter this model since we assume the rate-limiting step is these species traversing the synaptic cleft between neurons. **Evolution of tau**: The second term of Eq (**1a**) introduces a focal seeding event, at the entorhinal cortex (EC); i.e., where all misfolded pathology is produced, since there is historically accepted evidence that pathology begins in EC^1^. No other region is capable of producing misfolded pathology, but is able to further transmit it. **Evolution of Aβ**: Aβ spread is modeled similarly to tau, with diffusivity constant $\beta_{A\beta}$. Unlike tau, Aβ appears to have a diffuse production mediated by metabolism – this is encoded by the second term in Eq (**1b**). The production of misfolded amyloid is assumed to be proportional to both its baseline metabolism$\mathbf{m}$ measured via FDG-PET of healthy subjects, as well as the region’s pool of available Amyloid Precursor Protein (APP), $\mathbf{x}_{APP}$, from which $A\beta$ is cleaved.

**Driving function to model production of pathology at local sites**. Since onset of production is gradual, followed by eventual decline and plateauing, we govern the production terms by long-duration gamma-shaped driving functions $f_{A\beta}$ and $f_{\tau}$; for $A\beta$ and tau, respectively. This is modeled in this study as smooth but time-limited driving functions, in recognition of the fact that abrupt, discontinuous production functions are biophysically implausible, and that the production terms cannot continue *ad infinitum*, and must eventually decline in response to amyloid saturation, cell death and other factors. The function chosen for this purpose is one shaped like the Gamma curve given by $f_{\tau}\left( t \right)= \frac{1}{T_{\tau}}e^{-t/T_{\tau}}$, shown in **Figure S1**. The function depends on a single parameter $T_{\tau}$ that controls the duration of production of tau; analogously for amyloid. Trial and error resulted in the best choice of this parameter as $T_{\tau}=T_{A\beta}=5 years$. This function reproduces the expectation that at the beginning the rate of production is low, and steadily increases with increasing pathology. The rate of production then plateaus in response to countervailing forces in the brain. Finally, as neurodegeneration takes hold and amyloid and tau begin to be increasingly sequestered into plaques and tangles, the rate of production of soluble pathology begins to decline. It is stressed that the shape used here, Gamma function, is only one of many possibilities. Its choice was motivated by its smoothness, time boundedness, unimodal peak, and its ubiquity in mathematical modeling. This is clearly a design choice but not an essential one; any reasonable bounded driving function of sufficiently long duration will suffice.

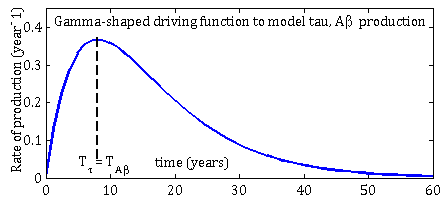

**Figure S1. Gamma-shaped production function for both tau and amyloid production**. Related to Experimental Procedures, Supplemental Experimental Procedures: **Note 3**. The y-axis represents the rate of production of each entity as a function of time, with t=0 at disease onset. This function gives low production in the beginning, hitting a peak in the middle of disease course, and thence declining.

**Modeling the network-mediated interaction between tau and amyloid**

Several possibilities of cross-interaction exist; they were listed and summarized in main Table 1. Here we specify those having the highest empirical or mechanistic support, and show how to encode them mathematically within the framework of network spread of both tau and amyloid.

2) **1-way interaction model: Amyloid affects tau aggregation**. A new cross-species interaction term was added via coupling coefficient $\gamma$ to simulate a potential role for amyloid to govern the rate of growth of misfolded tau ^16–18^:

Amyloid-facilitated tau aggregation: $\frac{d\mathbf{x}_{\boldsymbol{\tau}}(t)}{dt}= -\beta H\mathbf{x}_{\boldsymbol{\tau}}\left( t \right)+ {\alpha f}_{\tau}\left( t \right)\mathbf{e}_{EC}+\gamma(\mathbf{x}_{\boldsymbol{A\beta}}\left( t \right)\cdot\mathbf{x}_{\boldsymbol{\tau}}\left( t \right)\cdot(1-\mathbf{x}_{\boldsymbol{\tau}}\left( t \right)/K))$	(**2a**)

Note the last term has a logistic growth term of the form $x(1-\frac{x}{K})$. This nonlinear term is required to ensure that tau does not grow exponentially indefinitely, because realistic tau growth must be limited by overall capacity of healthy tau from which toxic tau can be cleaved. This capacity is controlled by the parameter K. The equation for amyloid evolution (**2b**) remains the same as (**1b**).

3) **1-way interaction model: Amyloid affects tau diffusion into the network**. In this model amyloid does not influence tau aggregation but increases the rate of tau diffusivity into the network in areas where amyloid is high and less diffusivity in areas where amyloid is low. This model simply modulates the scalar diffusivity constant in Eq (**1a**) by allowing it to be regionally-varying:

Amyloid-facilitated tau diffusion: $\frac{d\mathbf{x}_{\boldsymbol{\tau}}(t)}{dt}= -\beta\cdot diag\left\{ \mathbf{x}_{\boldsymbol{A\beta}}\left( t \right) \right\}\cdot H\mathbf{x}_{\boldsymbol{\tau}}\left( t \right)+ {\alpha f}_{\tau}\left( t \right)\mathbf{e}_{EC}$	(**3a**)

In this model tau in every region experiences a different net diffusivity, depending on the local level of amyloid. The equation for amyloid evolution (**3b**) remains the same as (**1b**).

4) **Remote amyloid to tau interaction model.** This model implements recent observations suggesting that amyloid affects tau aggregation, not locally but remotely, via long-range axonal projections i.e. the connectome^5,40^.

Remote amyloid-facilitated tau aggregation: $\frac{d\mathbf{x}_{\boldsymbol{\tau}}(t)}{dt}= -\beta H\mathbf{x}_{\boldsymbol{\tau}}\left( t \right)+ {\alpha f}_{\tau}\left( t \right)\mathbf{e}_{EC}+\gamma((C\mathbf{x}_{\boldsymbol{A\beta}}\left( t \right))\cdot\mathbf{x}_{\boldsymbol{\tau}}\left( t \right)\cdot(1-\mathbf{x}_{\boldsymbol{\tau}}\left( t \right)/K))$	(**4a**)

The equation for amyloid evolution (**4b**) remains the same as (**1b**).

5) **Tau to amyloid interaction model**. The reverse effect of tau on Aβ production^29^ does not have firm pathophysiological support ^16^. However, for completeness we also explored the alternative interaction via the coefficient $\eta$. The equation for tau evolution (**5a**) remains the same as (**1a**).

Tau-facilitated Aβ: $\frac{d\mathbf{x}_{\boldsymbol{A\beta}}(t)}{dt}= -\beta H\mathbf{x}_{\boldsymbol{A\beta}}\left( t \right)+ {\alpha f}_{A\beta}\left( t \right)(\mathbf{m}\cdot\mathbf{x}_{APP}) +\eta(\mathbf{x}_{\boldsymbol{\tau}}\left( t \right)\cdot\mathbf{x}_{\boldsymbol{A\beta}}\left( t \right)\cdot(1-\mathbf{x}_{\boldsymbol{A\beta}}\left( t \right)/K))$	(**5b**)

6) **Two-way interaction model**. Finally we included in our analyses a bidirectional model whereby both species influence each other. Hence Eq (**6a**) for tau evolution is the same as Eq (**2a**) and Eq (**6b**) for amyloid evolution is the same as Eq (**5b**).

**Modeling different modes of transmission**

7) **Proximity-based spread**. We simulated proximity-based spread using the same formulation as Eqns (**1a** and **1b**), but replacing the connectome-derived Laplacian with adjacency matrix computed from inter-regional Euclidean distance: $C_{dist}=\exp\left( -\frac{D}{\sigma} \right),$where $D=\{d_{ij}\}$ is the matrix of distance between regions i,j, and $\sigma$ is the standard deviation over all elements of D. Laplacian $H_{dist}$ was defined analogously to the connectome Laplacian $H$, and replaced that term in Eq (**1**). We also evaluated a Gaussian relationship instead of exponential, but the results were generally worse and were not further evaluated. We denote the full model by Eq (**7a**, **7b**).
8) **Fiber distance-driven spread**. We simulated spread along fiber projections using the same formulation as Eq (**1**), but replacing connectivity with fiber distance, as per $C_{fiberdist}=\exp\left( -\frac{L}{\sigma} \right),$where $L=\{l_{ij}\}$. Here $l_{ij}$is the average length of streamlines between regions i,j. Laplacian $H_{fiberdist}$ was defined analogously to the connectome Laplacian $H$. Denote the full model by Eq (**8a**, **8b**).

**Implementation**. The above theoretical models were numerically solved using MATAB’s **ode45()** solver, jointly for both amyloid and tau. The initial condition was set such that $x_{\tau, A\beta}(t=0)=0$; however note that the seeding and production of the two species are non-zero, via the terms $f(\cdot)$.

Note 2. Image processing and analysis of ADNI data

**Image acquisition**

High resolution T1-weighted MR images were available from the ADNI website from multi-site 3.0 T GE scanners using scanner-specific T1-weighted sagittal 3D MPRAGE sequences (8-channel coil, TR = 400 ms, TE = min full, flip-angle = 11°, slice thickness = 1.2 mm, resolution = 256 × 256 mm and FOV = 26 cm. AV45- and FDG-PET data are also available for download from on multiple instruments of varying resolution and following different platform-specific acquisition protocols. PET data in ADNI underwent standardized image pre-processing correction steps to increase data uniformity across the multicenter acquisitions. More detailed information on the different imaging protocols employed across ADNI sites and standardized image pre-processing steps for MRI and PET acquisitions can be found on the ADNI web site (<http://adni.loni.usc.edu/methods/> ). Demographic information is summarized in **Table S1**.

**Image processing**

The two datasets were processed independently with different processing streams, the former at Weill Cornell using image processing pipelines established at the Raj laboratory on the AAL atlas parcellation with 90 cerebral regions, and the latter using pipelines developed at the UCSF site on the 86-region Desikan atlas parcellation. Although both pipelines are derived from widely used tools (SPM, FSL, Freesurfer, Camino, etc) and have many common elements, their independent processing enables testing whether the proposed models are specific to our pipelines and atlas, or whether they are more generally applicable. Detailed processing steps are available elsewhere ^15,82^. Briefly, T1 images were normalized into MNI space and segmented using SPM8’s unified coregistration and segmentation scheme. GM was further parcellated into the 116 region of interest (ROI) AAL atlas ^83^ using the IBASPM package ^84^ within. The post-processed AV45-PET and AV1451-PET images were downloaded from ADNI; these images were already normalized to the same common space, scaled by cerebellar values and resliced to uniform voxel resolution. PET images from all normal controls were used to create an average image, which was then normalized to MNI space using SPM8’s linear and non-linear transformation. This transformation was then applied to all individual’s PET images, which were then resliced to the same resolution as the 90-region atlas at the Cornell site and to the 86-region atlas at the UCSF site. Mean AV1451- and AV45-PET signals were then calculated over each of the GM ROIs.

The raw imaging data were then converted into regional group statistics, following established approaches in neuroimaging. Groupwise, unpaired t-tests were computed between the MCI/AD groups and the normal controls for the AV45-PET and AV1451-PET signal averages in each GM region. T-statistics were calculated to maintain the convention that higher positive value signifies more pathology. Next, since subsequent analysis warrants positive and range-bound measures of regional effects, we scaled the t-statistic to have a mean of 1, and replaced all negative entries (that would represent artifactual hypertrophy) by 0.

**Connectomes.** Connectomes corresponding to the 90-region AAL parcellation were extracted from 69 healthy subjects, whose data was acquired under a previous study at Cornell, using a processing pipeline that is well established in the Raj laboratory; details can be found in ^82^. In short, T1 images were segmented and atlased using SPM8/IBASPM, orientation functions constructed using spherical deconvolution and probabilistic fiber tracking performed in subject’s DTI space at 2mm voxel resolution. The average connectivity across 69 subjects was computed for each region pair, to obtain a canonical healthy connectome C. A different canonical healthy connectome corresponding to the 86-region Desikan atlas was generated from the same 69-subject Cornell data, using the same Cornell pipeline.

Table S1. Sample size of the ADNI-3 diagnostic cohorts used in this study. AD: Alzheimer’s Disease patients, EMCI: early mild cognitive impairment, LMCI: late MCI, HC: healthy controls. These subjects represent the most complete set of subjects included in the ADNI-3 study as of 1/1/2021 who had all 3 imaging data (AV45-PET, AV1451-PET and MRI). The sample size listed here are for those subjects who had more than one tau PET scan.

|  | **Diagnosis** | **N** | **Age range** |
| --- | --- | --- | --- |
| ADNI 3 | AD | 68 | 73.8 (8.12) |
|  | EMCI | 137 | 72.1 (5.91) |
|  | LMCI | 75 | 71.9 (7.61) |
|  | HC | 251 | 74.10 (6.69) |

Note 3. Spatial Distribution of APP and MAPT

The rate of Aβ accumulation is modeled in our approach as the product of two starting distributions: a) the distribution of APP, the precursor protein from which Aβ is cleaved; and b) the distribution of regional metabolism, as measured from FDG-PET scans of healthy age-matched individuals in the ADNI cohort. APP regional patterns are not available from *in vivo* scans, hence a rough spatial pattern vector of APP was computed from publicly available human gene expression data and from post-mortem histology data. We relied on two studies to obtain the pattern of APP in the human brain. First, we used the Allen Brain Institute’s human brain gene expression data, from which the APP expression values were resolved on all available sample locations in 6 brains. This data, shown in **Figure S2** (left), strongly suggests that APP distribution in the brain is highly heterogeneous, with above-average expression in brainstem, subcortical and frontal cortices, but below average expression in posterior and cerebellar structures. This closely matches a previous histology study of post-mortem brains, which also found that APP, along with all the other Aβ-producing factors (PS1, BACE1, APP-CTFβ) are usually under-expressed in the posterior and parietal neocortex, as well as in cerebellum ^85^. For example, the expression of APOE, which plays a strong role in the processing of APP, is similarly distributed, with below average levels in posterior cortices and above-average levels in striatal and subcortical structures (**Figure S2**, right). The study by Shinohara et al ^85^ suggested that the numerical values of APP distribution in posterior cortices is approximately 50% compared to other cortices, especially in orbitofrontal cortex. Since APP levels from either our in-house or the study by Shinohara et al were not quantitatively available from the individual brain regions used in this study, we simply use a 2-level APP distribution, where all brain regions are assigned the APP level of 1, except the occipital regions, whose APP level is assigned 0.5. Note that the cerebellum is not part of this study, hence its APP level was not used. Some striatal areas, in particular putamen and caudate nucleus, show higher APP levels than the rest of the brain, as shown in ^85^. Therefore these two structures were assigned the relative score of 1.1.

Tau accumulation is not assigned a specific regional driver, however, since misfolded tau derived from physiologic tau, its concentration in turn is expected to be governed by the expression of microtubule associated protein tau (MAPT). MAPT expression is now quantitatively known in various parts of the brain, and in general is much lower in occipital and striatal regions ^86^. Hence we assign a rough map of MAPT with the value of 1 everywhere except occipital, putamen and caudate, where it is assigned 0.5.

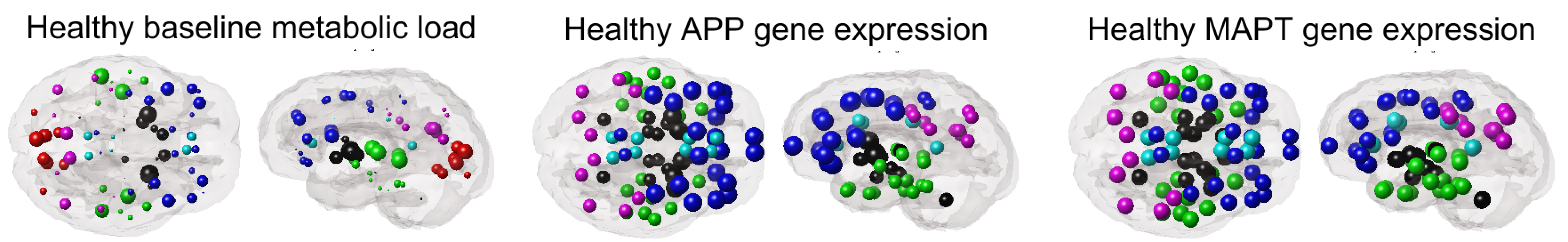

Figure S2. Spatial distribution in the human brain of factors driving the production of amyloid in the proposed model of Eq 1. These factors are: baseline glucose metabolic load and APP gene expression, and together they govern the spatial distribution of amyloid production, as per Eq 1. Recall that tau is deemed produced only at the seeding site, which is EC throughout the manuscript unless otherwise noted. To provide some context to the tau production aspect, on the right panel we show the spatial distribution of the MAPT gene. MAPT is regarded as the pool from which misfolded tau is cleaved. Cleary, MAPT does not show appreciable spatial concentration in the human brain, occurring diffusely throughout, with the exception of visual cortices. The metabolic load is the t-score of healthy FGD-PET SUVr in the cognitively normal subjects in ADNI. Spheres are located at the site of the gene expression sample, collected over 6 healthy young brain hemispheres by the Allen Institute. Sphere diameter is proportional to the z-score with respect to overall mean and standard deviation (+/- 2 std are shown here) and color-coded by lobe as in the main manuscript. Amongst cerebral regions, posterior and occipital regions have the least overall levels of APP and MAPT, while brainstem and subcortical structures have the highest levels. Frontal regions are generally above average in both.

Note 4. Global pathology burden predicted by the model

**Figure S3** shows the global accumulation of theoretical pathology over model time, evaluated at default parameters. All proteins increase over time, but amyloid-facilitated tau diverges dramatically from the non-facilitated tau at around t=15, mirroring main **Figure 3**. Clearly, while the “pure” tau model also captures empirical data, it does so far slower and achieves far less correlation strength than the facilitated version.

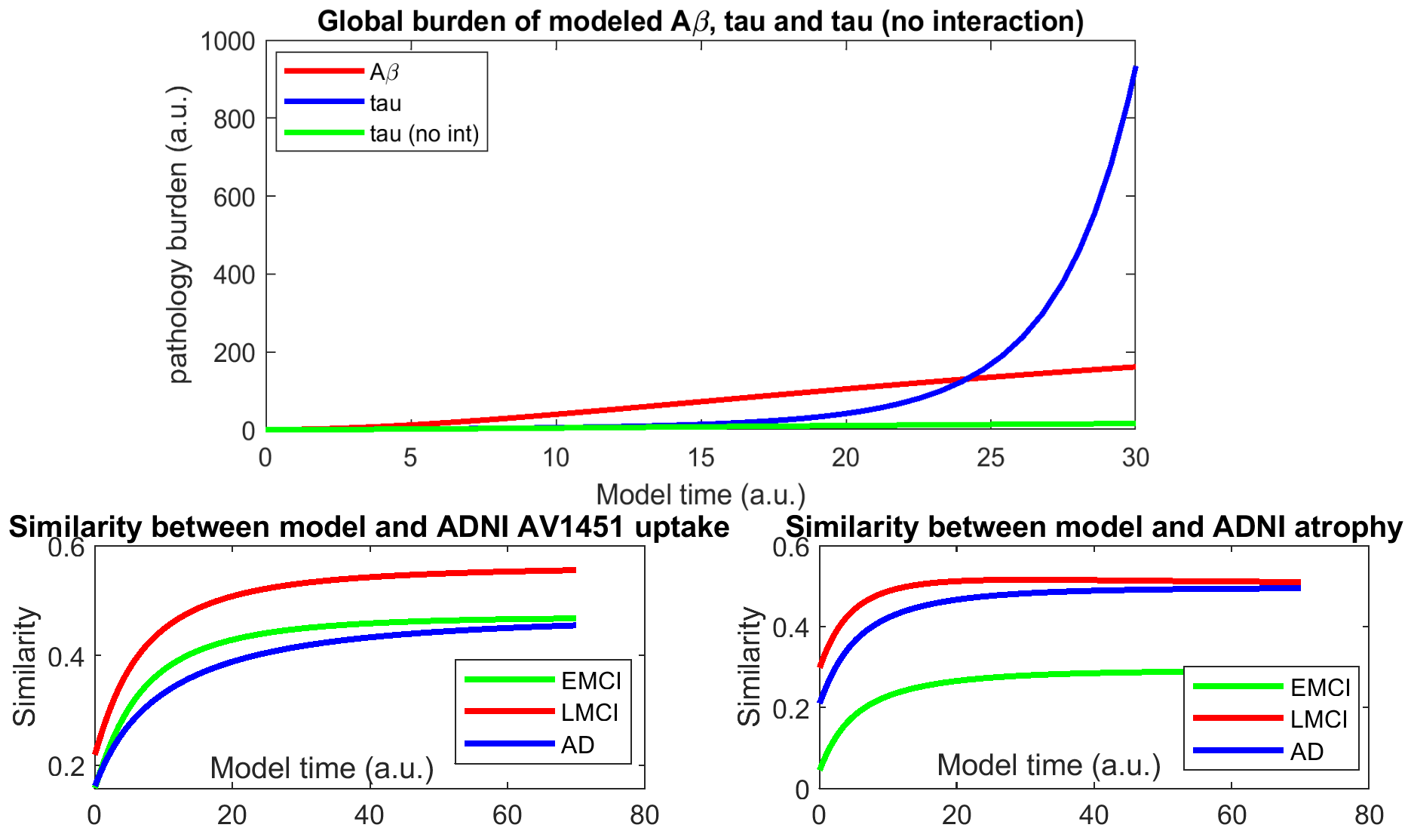

**Figure S3**. **Global burden of modeled pathology and the effect of amyloid facilitation. Top:** The evolution of global pathology burden as evaluated by numerical integration of Eqs (1) under default model parameters. **Bottom**: R-t curves showing the correlation between the non-facilitated tau model and empirical tau and atrophy data. Note the significantly longer time scale compared to **Figure 4**, and the lower peak R.

Note 5. Influence of Atlas

The two atlases used in this study, Desikan and AAL, give comparable results when the model is applied to each. **Figure S4** (left) shows the second eigenmode of the Laplacian from each connectome, with highly similar results. In both cases the highest values are in temporal cortices, which in this study was used to explain the early formation of tau fibrils in these regions. Similarly, the two atlases gave comparable distribution of empirical atrophy measured from MRI of the ADNI cohort, as shown in **Figure S4** (right). Note that neither the datasets, nor the atlases, nor the image processing pipeline used in constructing the two regional distributions are identical; this was described in **Supplemental Experimental Procedures: Note 1**. Thus, we conclude that the choice of connectome, atlas parcellation, processing pipeline or the specific stratification of the ADNI cohort are responsible for achieving the results shown in the main text.

A

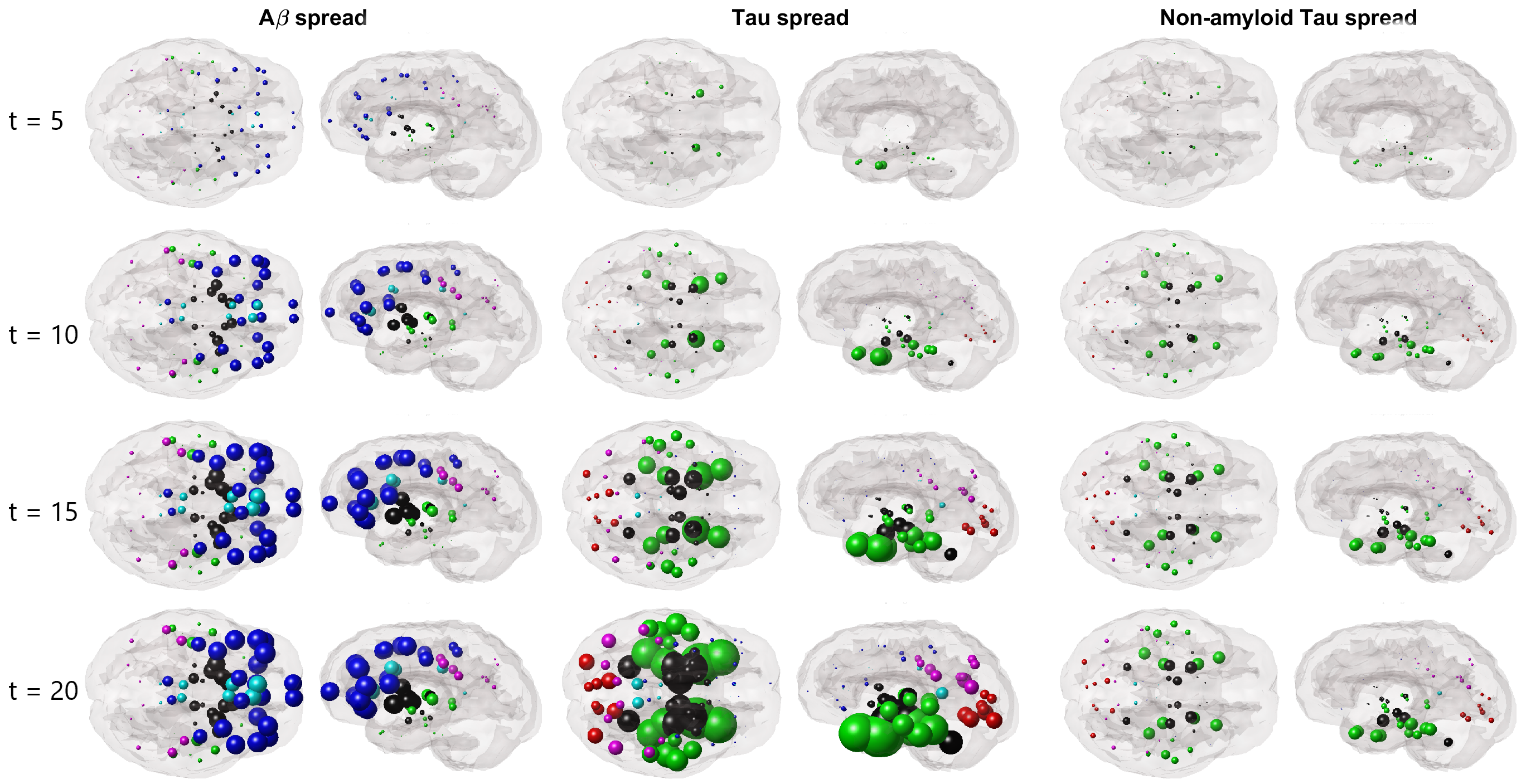

B

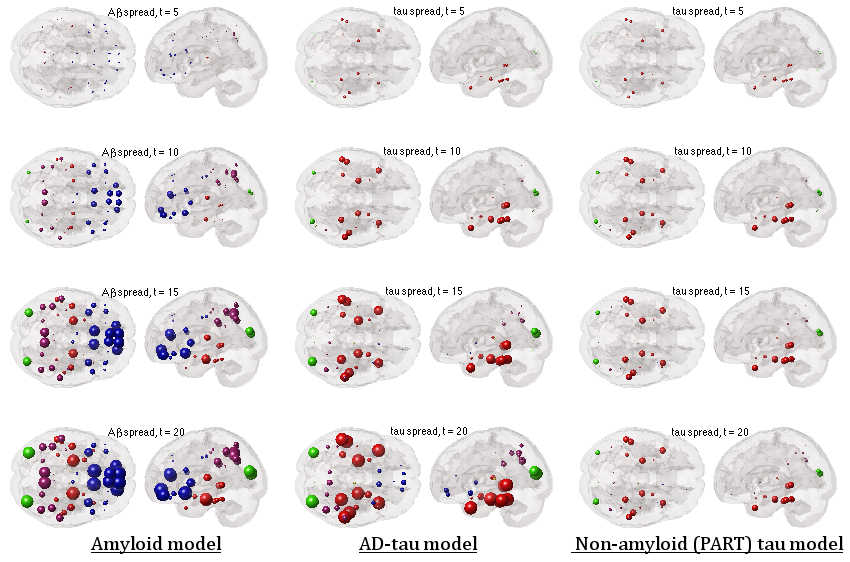

**Figure S4**. The evolution of model dynamics of amyloid and tau, evaluated by numerical integration of Eqs (1a,b), on the 90x90 healthy AAL connectome, under default model parameters. The first column shows the evolution of amyloid, the second column of amyloid-facilitated tau and the third column of tau only. Note the AAL atlas does not separate out entorhinal cortex, which is included within the parahippocampal gyrus. Spheres are color-coded by lobe, but color assignment is different from **Figure 4**.

Note 6. Parameter fitting for individual cases

The full models in Eqns (1,2) have 6 unknown global parameters: $\beta, \alpha, \gamma,\eta, T_{A\beta}, T_{\tau}, K$. Fortunately, the model is not overly sensitive to parameter choice, since all these parameters are related in one way or another to the time-scale of the neurodegenerative process, thereby primarily governing the scale of the time axis, not the overall behavior. We set the time-specific parameters $T_{A\beta},T_{\tau}$ at the outset heuristically over 20 trials to yield a temporal response that roughly matched the duration of Alzheimer pathology dissemination, i.e. 30 years ^87^. This yielded $T_{A\beta}=T_{\tau}=10 years$. The parameter *K*, which represents the capacity of overall pool of tau and amyloid, and thereby governs the logistic growth terms in the equations, was set at 5 for all subjects and simulations. The exact choice appears to not be important; similar model behavior was observed for other choices, like 10 or 20.

Other model parameters were optimized in two steps. First, starting with initial guesses of all parameters at 1.00, we employed brute-force grid search, over a range with 10 equally-spaced points per parameter. This yielded the parameter values shown in **Table 3**. We call this coarsely-optimized model specification the “default parameters”, which was used for assessing the model qualitatively and to understand its behavior over different seeding sites. Please note that the temporal units used here are shown as “years” for illustration only; since the model time as arbitrary units, so do the above rate constants.

Next, using the default parameters as starting guesses, we implemented a maximum a posteriori (MAP) estimator for the model, which jointly maximized a likelihood function as well as priors on each parameter. For this purpose we chose relatively shallow mutually-independent priors given by Gaussians around the initial guess of each parameter. The hyperparameter that weighs the prior term was obtained via trail and error, and set at 0.01; the method is not sensitive to the hyperparameter choice, since it is usually very small in order to not overly rely on the priors. We selected a likelihood function that is given by the strength of correlation to empirical data rather than more typical residual norm:

$$\Pr\left( x_{\tau}\left( t \right), x_{A\beta}\left( t \right)|\theta\right)\propto\exp\left( -R\left( x_{\tau}\left( t,\theta\right),x_{\tau}^{ADNI} \right) \right)\cdot\exp\left( -R\left( x_{A\beta}\left( t,\theta\right),x_{A\beta}^{ADNI} \right) \right)$$

$$t_{max}|\theta=\arg\max_{t} \prod_{i} \Pr\left( x_{\tau}\left( {t+\Delta t}_{i} \right), x_{A\beta}\left( t+\Delta t_{i} \right)|\theta\right)$$

where $R(\cdot, \cdot)$ is the correlation operator, $\theta$ is the collection of model parameters, and the model’s evolution is governed by these parameters via the ODE. Note that the fits employ all available longitudinal visits $t_{i}$, where by convention we define time gaps by ${\Delta t}_{i}=t_{i}-t_{1}$. The resulting MAP problem then becomes a non-linear cost minimization problem, which was solved using MATLAB’s fmincon() subroutine. Since fmincon() calls the ODE solver during each iteration, the process can become time consuming. Fortunately we found that using K=50 iterations was quite adequate; there was no discernible difference by using much higher number of iterations. Overall execution time for a single subject fitting ranged from 5 to 20 seconds.

**Table S2**: **Model parameters and defaults used in optimization.** Parameters set after coarse grid search (“coarsely optimized” or default parameters) and after further refinement using the Maximum a Posteriori estimation procedure (“Optimal values”), using the default parameters as initial guess. All parameters have units of $year^{-1}$, since they are rate parameters.

| Empirical data used for fitting | Coarsely optimized (default) parameters | Fitted parameters |
| --- | --- | --- |
|  | $\beta, \beta_{a}, \alpha, \alpha_{a}, \gamma$ | $\beta, \beta_{a}, \alpha, \alpha_{a}, \gamma$ |
| EMCI | 0.4, 1.00, 2.00, 2.00, 0.0 | 0.22, 0.86, 1.00, 1.33, 1.33 |
| LMCI | 1.00, 1.00, 2.00, 2.00, 0.20 | 0.28, 1.38, 0.99, 1.43, 1.44 |
| AD | 1.00, 1.00, 2.00, 2.00, 0.20 | 0.28, 0.97, 1.02, 1.22, 1.20 |

The other spread models (other than connectome-based spread) were optimized in the same fashion. Their optimal values did not change much or in interesting ways, likely due to the normalization of the adjacency matrices implicit in the definition of the Laplacian.

**Model fitting to individual subjects**. We applied the same MAP estimator to individual subjects, whether they had a single baseline scan or multiple longitudinal scans. In the latter case, the likelihood function above was evaluated jointly at all available longitudinal samples, keeping the time gap between them intact. Both amyloid and tau evolution occurs jointly in time in our model, hence in main **Figure 3** we report the correlation achieved by both tau and amyloid at the same time instant $t_{max}$, when the posterior probability peaks. It is possible to impose a delay between tau and amyloid onsets within our model, but we opted not to do so due to a lack of experimental data on that aspect.

Note 7. Permutation testing to demonstrate disease specificity

To determine whether the presented model displays AD-specificity and connectome-specificity, two permutation tests were devised:

A) 500 random permutations of the group AD atrophy pattern were obtained and repeatedly tested against the network model (**Figure S5-A)**. If the model is not specific to AD, a significant proportion of these tests should be positive. In fact, randomly permuted R cluster near zero; hence reported R values are highly significant compared to this “null” distribution ($p<{10}^{-3}$ for all groups). To simplify presentation and avoid excessive computational burden, in these simulations only linear and cross-product terms were retained in the models; this choice does not alter the overall system behavior while making computations dramatically faster.

B) The above null models tend to destroy the correlation structure in regional tau and atrophy. To avoid this and to test whether the true anatomic network structure is necessary to replicate our results, we performed 500 random permutations of the connectome itself, and correlated the (refitted) model seeded at EC against empirical (unchanging) regional pattern of tau-PET. Each connectome realization randomly scrambled the upper triangular portion of the true human connectome, and generated a symmetric matrix from that. Thus, these connectomes preserve actual connectivity values, but randomize network topology. If the model is not specific to the connectome, a significant proportion of these tests should be positive. The “null” R distributions shown in **Figure S5-B** are significantly smaller compared to the true reported R ($p<{10}^{-3}$ for all groups). The results for atrophy and amyloid are largely the same and not reported here. Note there is a hard lower limit in each case, which reflect the correlation between EC seed and regional tau prior to any network spread, regardless of the connectome used.

Taken together, these results confirm that the proposed model only recapitulates empirical regional distributions when it is applied in the correct region order and to the correct human connectome.

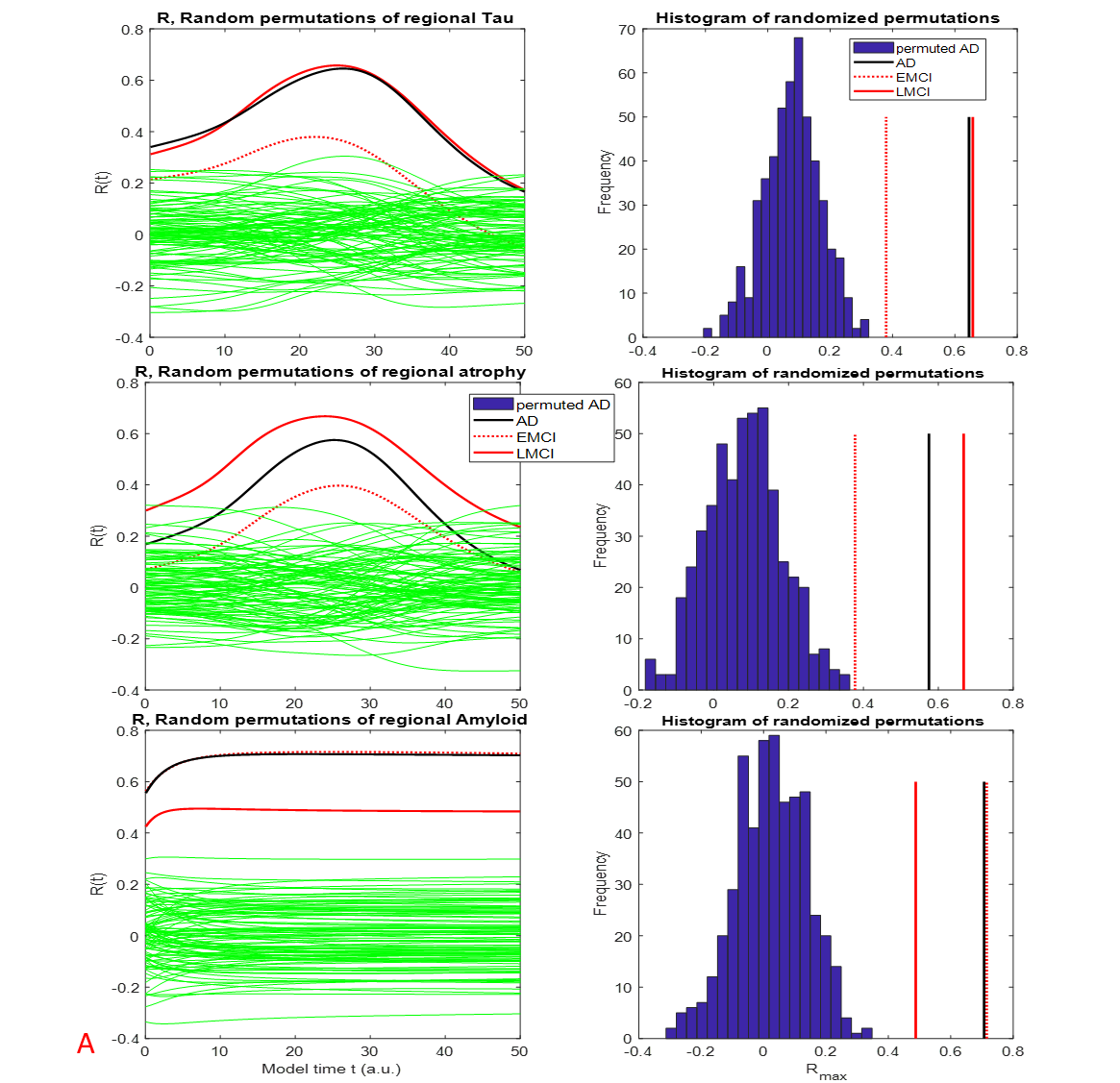

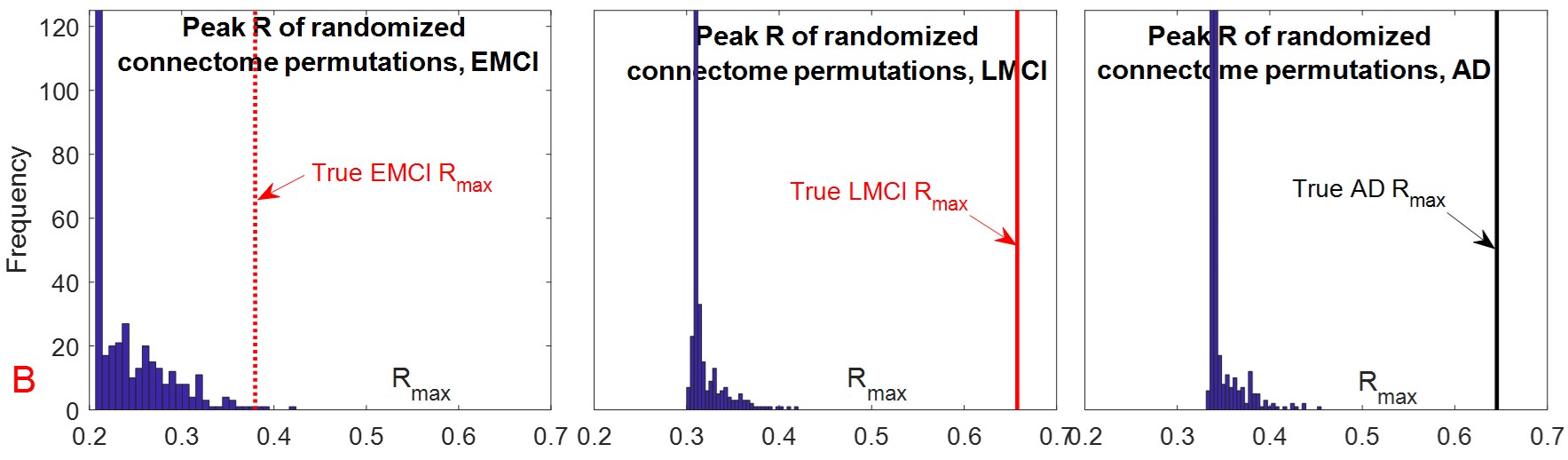

**Figure S5**. **Permutation testing**. Pearson’s R of correlation strength between proposed model in Eq (1a,b) solved via numerical integration, and empirical regional patterns of each diagnosis group (EMCI, LMCI, AD) in the ADNI cohort. **A**: Modeled vs empirical tau (top row), atrophy (middle) and Aβ data (bottom) are shown alongside the R-t curves corresponding to 500 random permutations of the regional patterns of the AD cohort. The random permutations lead to peak correlations with the model that are centered near 0, suggesting that the actual topography of the regional values is critical to the results we have reported. Histograms of peak R from random permutations are shown in right column, alongside R reported for empirical data, indicating that the latter are highly significant under permutation testing. **B**: Histograms of peak R against reginal tau from random permutations of the connectome, alongside R reported for true connectome, indicating that the latter are highly significant under permutation testing.

Note 8. Spearman correlations, compared to Pearson in Figure S5

The previous figure reported the agreement between the model and empirical regional distribution of biomarkers using Pearson correlation. It is now demonstrated that the results are similar when using a non-parametric correlation statistic, i.e. Spearman rank correlation. Spearman does not require that the data being compared are drawn from Gaussian distributions, hence it may be appropriate in the current context. **Figure S6** shows that the results using Spearman are generally similar to the main Pearson results in the main text but with lower significance. Pearson was preferred in this study because it is not as non-linear as Spearman, which is overly sensitive to low effect sizes, which generally pertain to brain regions that do not participate in AD pathology. These noisy values are largely irrelevant in the current context, yet Spearman rank is overly sensitive to these values. However, the fact that both Spearman and Pearson are in rough agreement gives additional confidence that the data reported in the main study are robust.

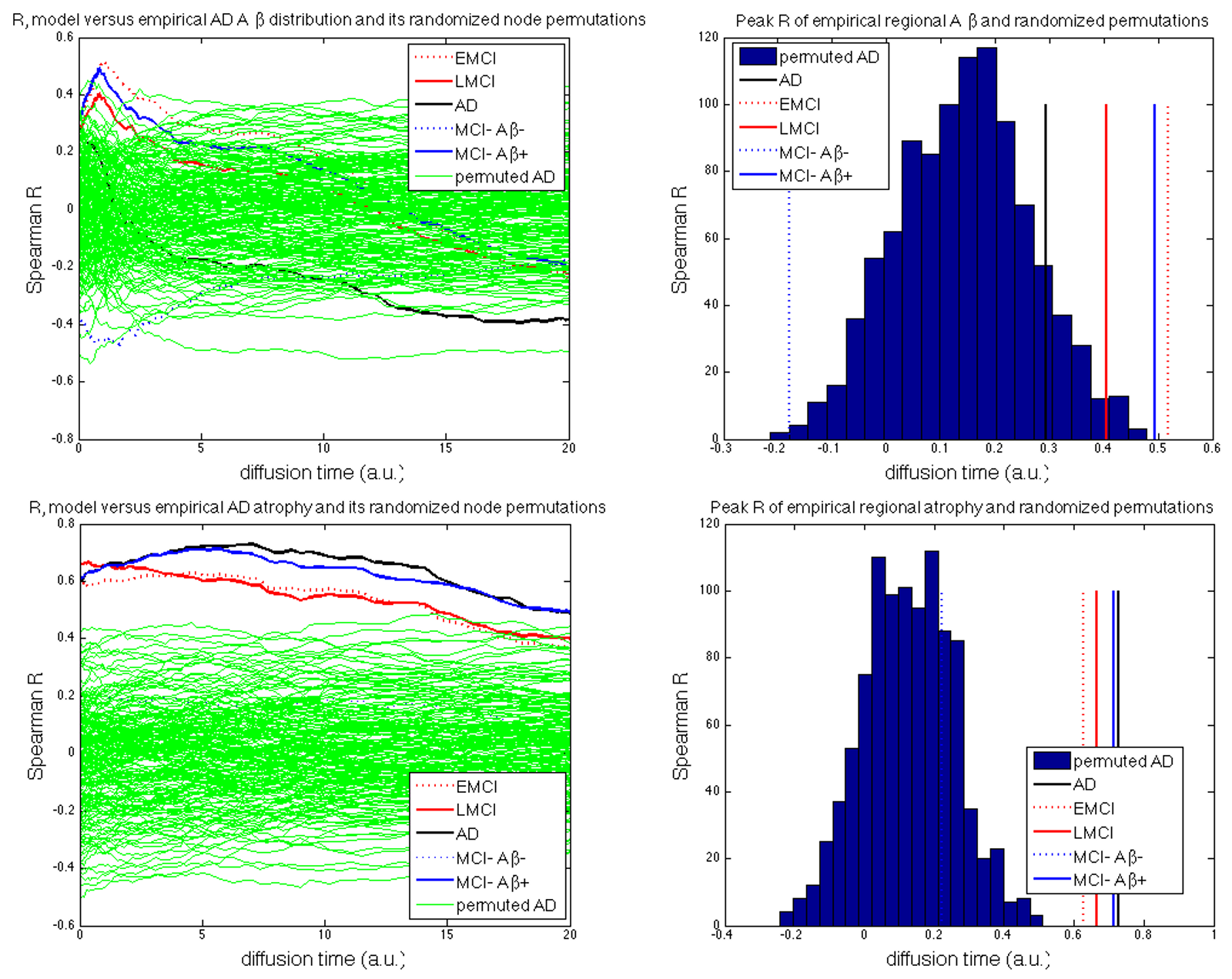

**Figure S6. Spearman correlation results of default model against empirical regional statistics**. Compared to Pearson correlation curves in main text, these results are very similar, if noisier due to non-linearities inherent in Spearman rank calculations. Both atrophy and amyloid patterns of the theoretical model are specific to AD phenotypes, and do not correlate well with either random permutations or MCI data for amyloid-negative subjects (dotted blue curves).

Note 9: Repeated seeding experiments to assess alternative seeding sites

Other than the canonical Entorhinal cortex (EC) seeding site, we also assessed other regions’ seeding plausibility. For this, we repeatedly simulated the joint model seeded from each region bilaterally. For each seed region, peak Pearson’s R between model and ADNI tau PET data were shown in **Figure 5**. EC is amongst the best overall cortical seeding site, while hippocampus (HP) is the best subcortical site. Other prominent seeding sites include Inferotemporal cortex (IT), Parahippocampal gyrus (PHP) and Fusiform gyrus (Fus). The evolution of pathology from these non-EC sites are shown below in **Figures S8** and **S9**, for HP-seeding and IT-seeding, respectively. They remain substantially similar to main **Figure 4**; however a notable difference from EC seeding is that HP seeding leads to higher involvement of medial temporal and subcortical structures. Amongst the above 5 regions, EC is unique in giving consistently one of the highest seeding likelihood despite having less levels of empirical tau deposition than others.

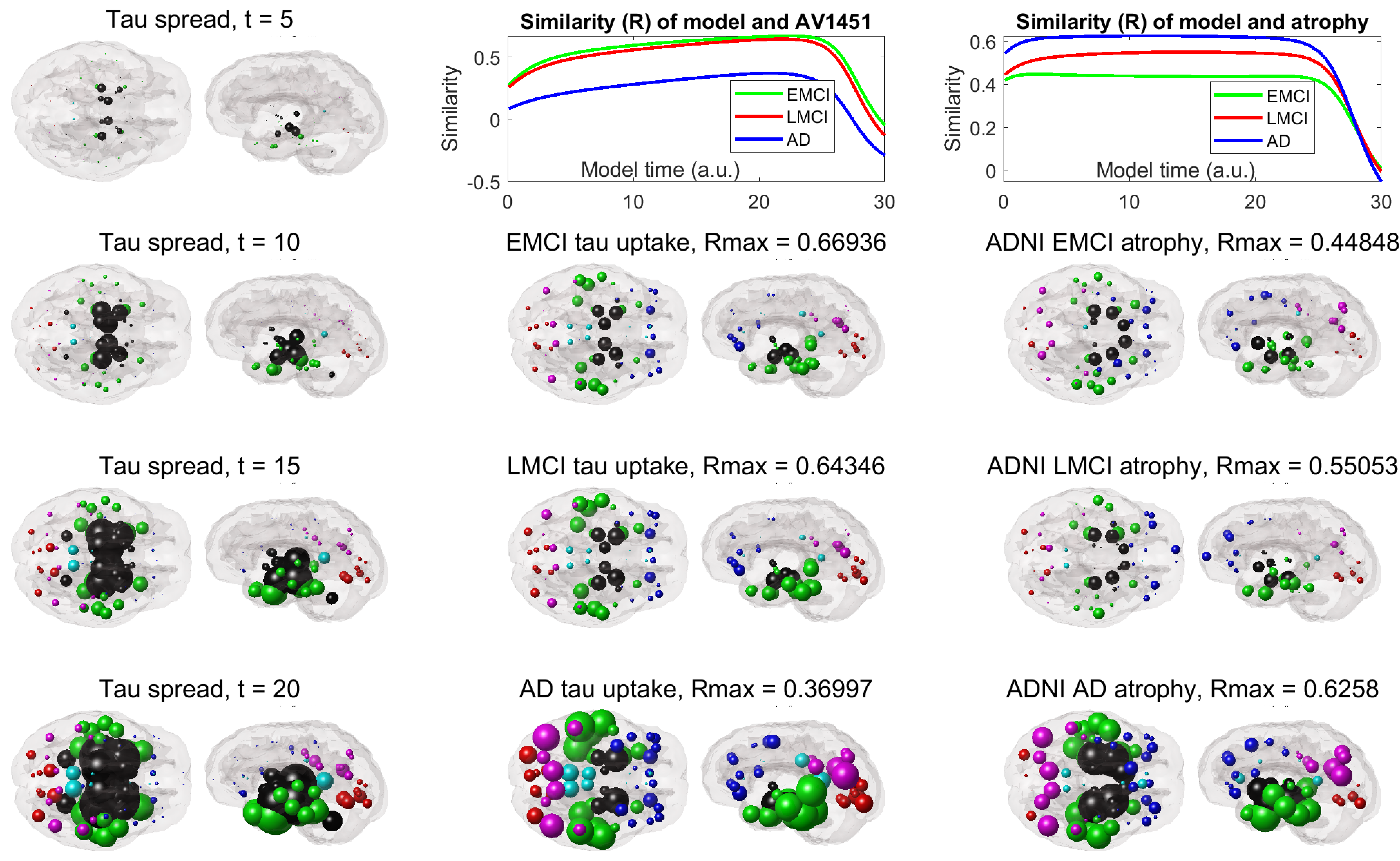

**Figure S7**. **Evolution of default model’s tau after seeding the Hippocampus, compared against empirical data.** The first column shows the modeled evolution of amyloid-facilitated tau, while the second column shows cross-sectional empirical tau-PET SUVr patterns from all 3 diagnostic groups. The top panel shows the behavior of peak R against model time t. The match between the model and empirical tau are strongest for E/LMCI and weakest for AD. The right-most column shows results for MRI-derived atrophy. The match between the model and empirical data are strongest for AD and weakest for EMCI. Peak R is highly significant in all cases. These results are similar to the base case presented in the main manuscript, **Figure 4**, which had EC seeding of tau. The key difference is that the baseline correlation (left-most end of the R-t curves shown in top row) start off higher than does EC – this is likely due to the stronger baseline levels of both tau and atrophy in hippocampus compared to EC. Another notable difference from EC seeding is that HP seeding lead to higher involvement of medial temporal and subcortical structures (denoted in black).

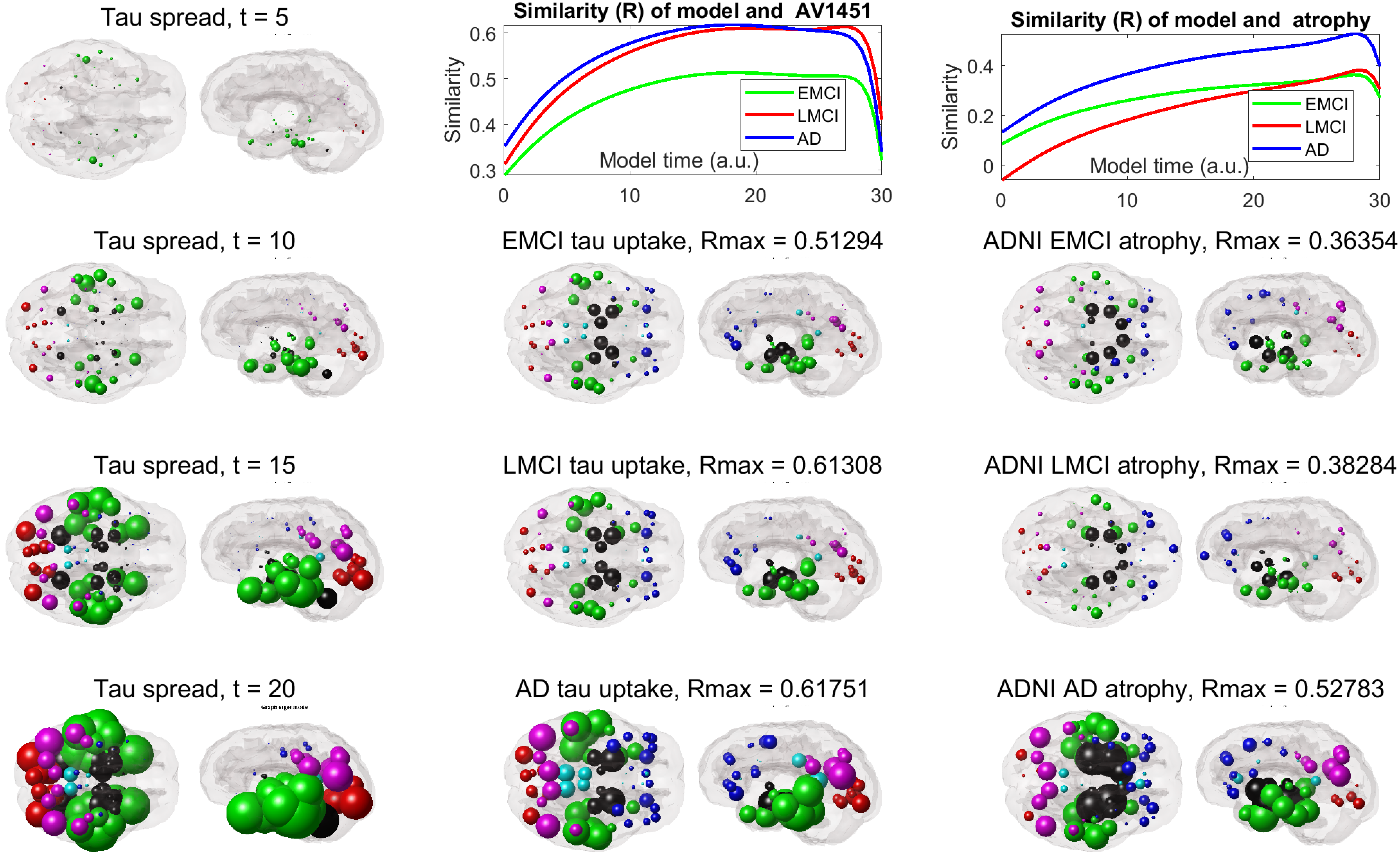

**Figure S8**. **Evolution of default model tau after seeding the Inferotemporal Cortex, compared against empirical data.** The first column shows the modeled evolution of amyloid-facilitated tau, while the second column shows cross-sectional empirical tau-PET SUVr patterns from all 3 diagnostic groups. The top panel shows the behavior of peak R against model time t. The match between the model and empirical tau are strongest for LMCI and AD, and weakest for EMCI. The right-most column shows results for MRI-derived atrophy. The match between the model and empirical data are strongest for AD and weakest for E/LMCI. Peak R is highly significant in all cases. These results are similar to the base case presented in the main manuscript, **Figure 4**, which had EC seeding of tau. The key difference is that the baseline correlation of tau (left-most end of the R-t curves shown in top row) start off higher than does EC – this is likely due to the stronger baseline levels of tau (but not atrophy) in IT compared to EC.

**Table S3**: **Model comparison using** **Fisher’s R-to-z transformed t-test results pertaining to main data in Table 2**. Six theoretical interaction models’ evaluations are presented: the No-interaction model, without the interaction term ($\gamma=0$); the 1-way interaction, tau🡪$A\beta$ aggregation, whereby tau affects amyloid but not vice versa; the 1-way interaction, $A\beta$🡪tau diffusion model, whereby amyloid influences the rate of diffusion of tau but not vice versa; the 1-way interaction, $A\beta$🡪tau aggregation model, whereby amyloid affects tau but not vice versa; the 1-way interaction, $C\cdot A\beta$🡪tau aggregation model, whereby distant amyloid affects tau but not vice versa; and the 2-way interaction $A\beta$🡨🡪tau aggregation model where both tau and amyloid affect each other via $\gamma$. Fisher’s R-to-z transformation was applied, and the *1-sided* p-value of significant differences between each model-pair was evaluated; hence only improvements from base model are assessed. P-values that are smaller than the significance threshold (p<0.05) are highlighted in bold red font, and indicate comparisons where one model is significantly different than another. For conciseness, Fisher comparisons are shown only for model-pairs “closest” to each other.

| **Model A:** | 1-way interaction tau🡪$A\beta$ aggregation | 1-way interaction $A\beta$🡪 tau diffusion | 1-way interaction $A\beta$🡪 tau aggregation | 1-way (remote) interaction $C\cdot A\beta$🡪tau aggregation | 2-way interaction $A\beta$🡨🡪tau aggregation |
| --- | --- | --- | --- | --- | --- |
| **Model B:** | No-interaction | No-interaction | No-interaction | 1-way interaction $A\beta$🡪 tau aggregation | 1-way interaction $A\beta$🡪 tau aggregation |
| EMCI Aβ | p = 1 | p = 1 | p = 1 | p = 1 | p = 1 |
| LMCI Aβ | p = 1 | p = 1 | p = 1 | p = 1 | p = 1 |
| AD Aβ | p = 1 | p = 1 | p = 1 | p = 1 | p = 1 |
| EMCI tau | p = 1 | p = 1 | **p < 0.01** | p = 1 | p = 1 |
| LMCI tau | p = 1 | p = 1 | **p < 0.01** | p = 1 | p = 1 |
| AD tau | p = 1 | p = 1 | **p < 0.01** | p = 1 | p = 1 |
| EMCI atr | p = 1 | p = 1 | p = 1 | p = 1 | p = 1 |
| LMCI atr | p = 1 | p = 1 | p = 1 | p = 1 | p = 1 |
| AD atr | **p < 0.01** | p = 1 | **p < 0.01** | p = 1 | p = 1 |
